# Spatially global effects of feature-based attention in functional subdivisions of human subcortical nuclei

**DOI:** 10.1101/2024.10.26.620447

**Authors:** Weiru Lin, Yan-Yu Zhang, Xilin Zhang, Fang Fang, Chencan Qian, Peng Zhang

**Affiliations:** State Key Laboratory of Brain and Cognitive Science, Institute of Biophysics, Chinese Academy of Sciences, Beijing, 100101, China; University of Chinese Academy of Sciences, Beijing, 100049, China; School of Psychology, Shanghai Jiao Tong University, Shanghai, 200030, China; Key Laboratory of Brain, Cognition and Education Sciences, Ministry of Education, South China Normal University, Guangzhou, Guangdong, 510631, China; School of Psychology, Center for Studies of Psychological Application, and Guangdong Provincial Key Laboratory of Mental Health and Cognitive Science, South China Normal University, Guangzhou, Guangdong, 510631, China; School of Psychological and Cognitive Sciences and Beijing Key Laboratory of Behavior and Mental Health, Peking University, Beijing, 100871, China; IDG/McGovern Institute for Brain Research, Peking University, Beijing, 100871, China; Peking-Tsinghua Center for Life Sciences, Peking University, Beijing, 100871, China; Key Laboratory of Machine Perception (Ministry of Education), Peking University, Beijing, 100871, China

## Abstract

Attention can prioritize the processing of non-spatial features throughout the visual field. While human subcortical nuclei are known to play important roles in spatial attention, the subcortical mechanisms of feature-based attention remain elusive. Using high-resolution 7T fMRI, we investigated the spatially global effects of color-based attention in functional subdivisions of human subcortical nuclei. Paying attention to a color matching the unattended stimulus across the visual field selectively modulated BOLD signals in the parvocellular (P) layers of the lateral geniculate nucleus (LGN) of the thalamus, enhancing the response to the unattended stimulus while reducing the response to the attended stimulus. The spatially global effects of color-based attention were also found in the deeper layers of the superior colliculus (SC), early visual cortices, and the intraparietal sulcus (IPS) of the parietal lobe. Effective connectivity analyses further revealed enhanced feedforward and feedback connectivity between LGN and V1, along with top-down modulation from IPS through the SC and ventral pulvinar. Finally, attended color can be decoded from multi-voxel response patterns in frontoparietal regions. These findings demonstrate that color-based attention selectively modulates color processing in P subdivisions of the LGN of the thalamus throughout the visual field, controlled by top-down signals from the parietal cortex through the SC and pulvinar.

**Highlights:**

- Color-based attention selectively modulates parvocellular activity in the LGN throughout the visual field.
- Spatially global effect of feature-based attention significantly modulates activity in the deeper layers of the SC.
- Feature-based attention enhances both feedforward and feedback connectivity between the LGN and V1.
- Top-down signals from IPS through the SC and pulvinar controls feature-based attention.

## Introduction

Selective attention allows our brain to process behaviorally relevant information in cluttered visual scenes. Attention can be directed to spatial location, or to non-spatial features such as color or motion direction. In feature-based attention, items shared the same feature as the attended target can be prioritized throughout the visual field, independent with the spatial location of the visual items (Liu & Hou, 2011; Scolari et al., 2014; A. L. White & Carrasco, 2011). While the neural mechanisms of spatial attention has been extensively studied in both cortical and subcortical regions (Carrasco, 2011; Nobre & Kastner, 2014), much less is known about the neural mechanisms of feature-based attention, especially in subcortical regions.

Paying attention to the non-spatial feature of a stimulus selectively activates sensory cortex specialized for processing the attended feature dimension, such as motion-sensitive area MT or color-sensitive area V4 (Chawla et al., 1999; Corbetta et al., 1990). Within these brain regions, neuronal activity is either enhanced or suppressed depending on the similarity of the attended feature value and the neuronal feature tuning, even when the receptive field of the neuron is located far outside the focus of spatial attention (Martinez-Trujillo & Treue, 2004; Treue & Martinez Trujillo, 1999). The spatially global effect of feature-based attention has also been demonstrated in neuroimaging studies (Saenz et al., 2002; Serences & Boynton, 2007; W. Zhang & Luck, 2009; X. Zhang et al., 2018), further supporting the feature-similarity gain model of attention (Treue & Martinez Trujillo, 1999). Beyond the visual cortex, attended features can be decoded from multivoxel patterns in the frontoparietal regions (Liu et al., 2011). Causal evidence from electrophysiological study in monkeys (Bichot et al., 2015) and effective fMRI connectivity in humans (X. Zhang et al., 2018) support the prefrontal cortex as a source for controlling feature-based attention. However, a recent study showed that the attentional gain modulation of PFC neurons is independent with the feature tuning of neurons in a feature-based attention task (Stalter et al., 2021). Thus, the neural mechanisms of feature-based attention in frontroparietal cortex remains unclear.

In subcortical regions, neural mechanisms of spatial attention have been well studied in the lateral geniculate nucleus (LGN), thalamic reticulate nucleus (TRN) and pulvinar of the thalamus (Halassa & Kastner, 2017; Saalmann et al., 2012; Saalmann & Kastner, 2009), and in the superior colliculus (SC) of the brainstem (Krauzlis et al., 2013). LGN activity can be strongly modulated by top-down spatial attention (McAlonan et al., 2008; O’Connor et al., 2002; Schneider & Kastner, 2009), with early modulation by the TRN and late modulation likely through cortical feedbacks (McAlonan et al., 2006, 2008). The pulvinar plays a critical role in the control of spatial attention (Petersen et al., 1987) and regulates information transmission between cortical areas at the attended location (Saalmann et al., 2012; Zhou et al., 2016). The superficial and deeper layers of the SC have been proposed to encode the saliency and priority maps of attention, respectively (Katyal et al., 2010; Katyal & Ress, 2014; B. J. White et al., 2017). While much evidence supports the important roles of these subcortical nuclei in spatial attention, little is known about the subcortical mechanisms underlying feature-based attention. The only human fMRI study about feature-based attention showed stronger BOLD signals in the LGN and pulvinar when participants attended to motion compared to color in spatially overlapping fields of moving and colored dots (Schneider, 2011). However, the spatially global effects of feature-based attention in functional subdivisions of the subcortical nuclei have not been convincingly demonstrated.

After the retina, the LGN is the earliest stage in the visual pathway that processes feature-selective information. It consists of two magnocellular (M) layers and four parvocellular (P) layers in the ventral and dorsal parts of the nucleus, respectively. Neurons in the P layers prefer color, high spatial frequency and low temporal frequency stimuli, whereas neurons in the M layers are sensitive to motion, and luminance contrast at low spatial frequency and high temporal frequency (Derrington et al., 1984; Derrington & Lennie, 1984). Feature-selective neurons have also been found in the visual pulvinar (Casanova et al., 2001; Felsten et al., 1983), which consists of several functional subdivisions with distinct connectivity patterns with cortical regions. The ventrolateral pulvinar (vlPul) is interconnected with the early visual cortex and the ventral visual stream, whereas the ventromedial pulvinar (vmPul) receives projection from the SC and connects to the dorsal visual stream (Arcaro et al., 2015; Bridge et al., 2016; Kaas & Lyon, 2007).

Therefore, it is possible that feature-based attention could modulate feature-selective processing in functional subdivisions of the visual thalamus, controlled by cortical and subcortical attention network regions. Recent studies showed that high-resolution fMRI is capable to resolve fine-scale activity in functional subdivisions of human subcortical nuclei (De Martino et al., 2013; Denison et al., 2014; Y. Qian et al., 2020; Schneider et al., 2004; P. Zhang et al., 2015). Thus, to investigate the neural mechanism of feature-based attention in human subcortex and its relationship with cortical regions, we used high-resolution fMRI (1.5 mm isotropic voxels) at 7 Tesla to investigate the spatially global effects of color-based attention in the functional subdivisions of human subcortical visual nuclei (i.e., LGN, Pulvinar and SC), early visual cortices, and frontoparietal attention networks.

## Results

We carefully calibrated visual stimuli to isolate L/M cone opponent color processing in the early visual areas, particularly the P layers of the LGN (Derrington et al., 1984). In the behavioral experiment, we investigated whether paying attention to the same compared to different colors across the visual field could improve color discrimination performance in dual tasks (Saenz et al., 2003). In the fMRI experiments, we investigated whether the attended color matching the color of the unattended stimulus across the visual field could significantly modulate BOLD signals in the brain regions of interest (Experiment 1, Exp. 1), and whether the attended color could be decoded from brain activity to spatially overlapping red/green stimuli presented at attended and unattended locations (Experiment 2, Exp. 2).

### Attention to the same color in both hemifields improves color discrimination performance in dual tasks

In the behavioral experiment (Fig. 1a), participants performed two color discrimination tasks simultaneously at two different locations across the visual field. At the beginning of each trial, two colored dots next to the fixation point indicated the colors of target dots to be attended. After pressing a button, two spatially overlapping fields of red and green dots were presented on each side of fixation in two 1000-ms intervals, separated by a 100-ms fixation period. Participants paid attention to the cued dots, and responded in which interval the stimulus had higher red/green cone contrast. Feedback was given after responses. The result showed significantly better performance when the attended colors were the same compared to different across the visual field (Fig. 1b, t(24) = 9.425, p < 0.001, Cohen’s d = 1.202). This replicates the previous findings by Saenz and colleagues (Saenz et al., 2003), suggesting that feature-based attention can facilitate the processing of the attended feature throughout the visual field.

**Figure 1.**
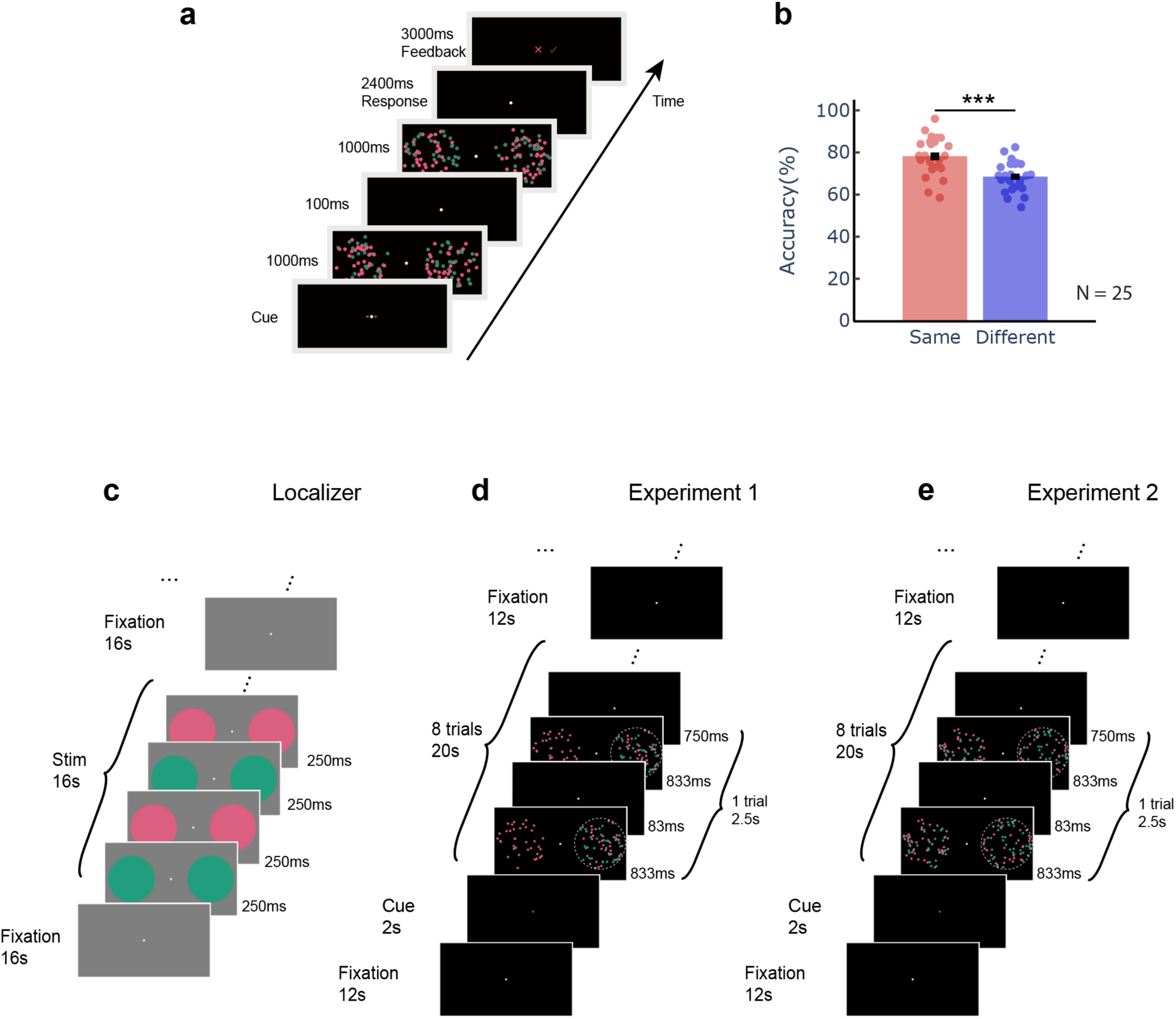
Schematic diagram of stimuli and procedures for the behavioral and fMRI experiments. **a.** Stimuli and procedure for one trial in the behavioral experiment. **b.** Color discrimination performance in behavioral experiment. **c.** Stimuli and procedure for the functional localizer in the fMRI session. Uniform red and green disks in L-M cone contrast flickered at 2 Hz for 16 s, alternating with 16-s fixation on gray background. **d.** Stimuli and procedure for Exp. 1. Participants attended to the cued dots at the cued location (indicated by the white dotted circle, which was not shown in the actual experiment), while ignoring the dots on the opposite side. 20-s task blocks alternated with 12-s fixation periods. **e.** Stimuli and procedure in Exp. 2 were the same as in Exp. 1, except that both red and green dots were presented at the unattended location. Error bars represent standard errors of the mean (SEM) across participants.

### Color similarity between attended and unattended stimuli across the visual field selectively modulates P layer activity in the LGN

In the fMRI session, isoluminant L-M cone-contrast defined red/green disk flickers were used as a chromatic functional localizer to selectively activate the P layers of the LGN (Fig. 1c). In Exp. 1 (Fig. 1d), participants attended to and discriminated the cone contrast of one of the two spatially overlapping red/green dot fields while ignoring the colored dots on the opposite side of fixation. The ignored dot fields were presented in either the same or different color as the attended dots. Behavioral performance didn’t show a significant difference between the same (mean ± std: 73.48% ± 8.75%) and different (73.44% ± 9.44%) conditions (t(24) = 0.030, p = 0.976, BF_10_ = 0.211). In Exp. 2 (Fig. 1e), spatially overlapping red and green dots were presented in both attended and unattended visual fields. Participants performed the same task as in Exp. 1 (71.42% ± 8.66%).

The anatomically delineated LGN mask was divided into M and P subdivisions based on the ventromedial-to-dorsolateral laminar organization of the human LGN, at a volume ratio of 1:4 (Fig. 2a and Fig. S1) (Andrews et al., 1997). The chromatic localizer activated the P subdivision in the dorsolateral LGN (Fig. 1b), whereas the colored moving dots presented on black background strongly activated both P and M subdivisions (Fig. 2c and 2d). Fig. 2c shows the activation maps in Exp. 1 when the attended color and the color of unattended stimulus in the opposite visual field were the same or different, and the response difference between the two conditions. The differential map between the same and different conditions shows a stronger activation in the P subdivisions of the LGN contralateral to the unattended visual field.

**Figure 2.**
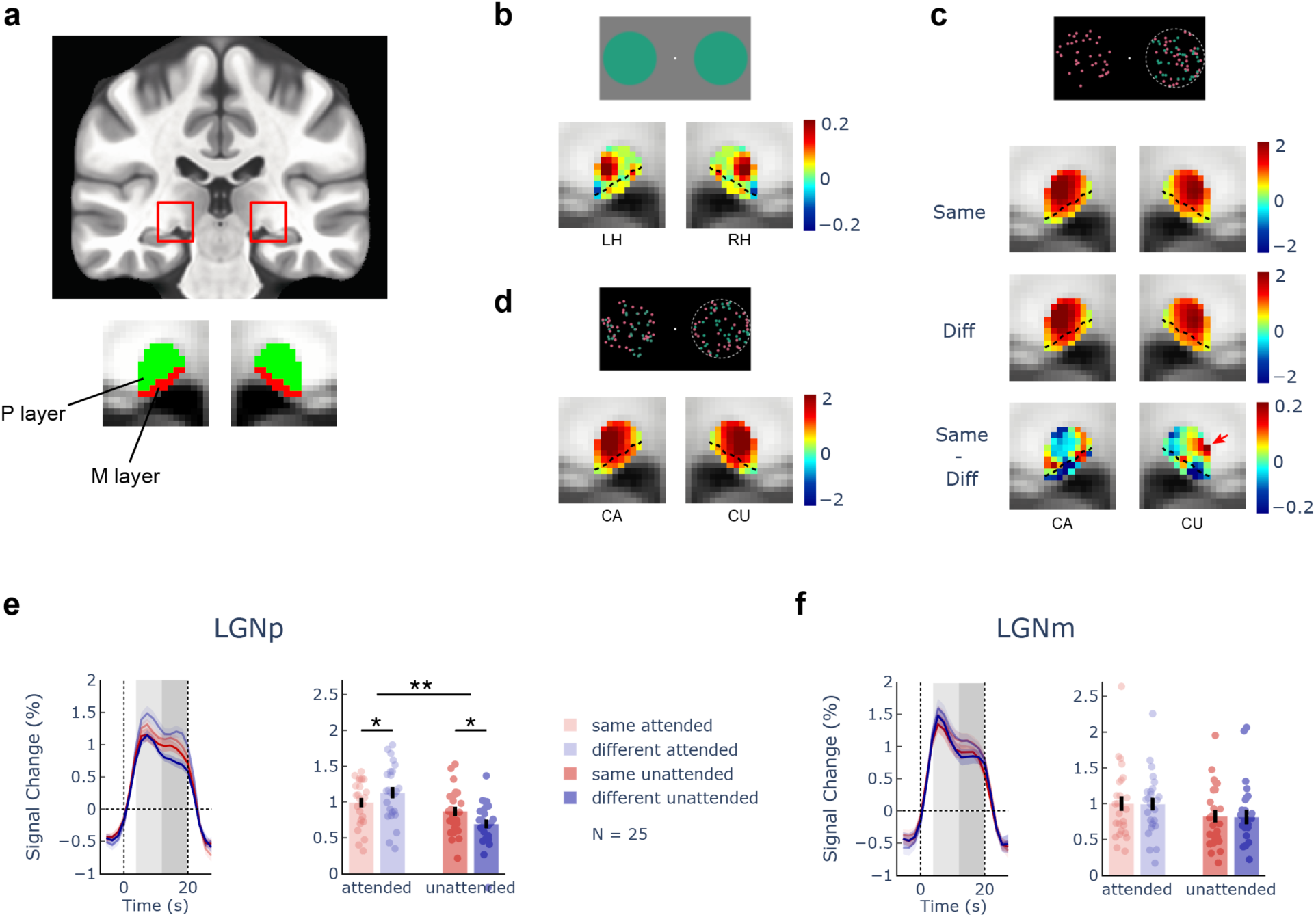
BOLD activation maps and ROI-averaged responses in functional subdivisions of the LGN. (a) The location of LGN in MNI space and the definition of M and P subdivisions in a coronal section. (b) (c) (d) show the activation maps to stimuli in the chromatic localizer, Exp. 1 and Exp. 2, respectively. The red arrow in (c) highlights the activation in the P subdivision to the spatially global effect of color-based attention. Colorbars indicate the percent signal change of BOLD signals. LH, left hemisphere. RH, right hemisphere. CA, contralateral to the attended location. CU contralateral to the unattended location. (e) (f) are ROI-averaged responses in LGNp and LGNm. Light and dark shaded areas in the timecourse plots indicate the transient and sustained response period, respectively. Shaded areas and error bars in the bar graph indicate SEM across participants. *: p < 0.05, **: p < 0.01.

To further investigate the spatially global effects of color-based attention in Exp. 1, we analyzed the ROI-averaged BOLD response to visually responsive voxels in the M and P subdivisions. P-biased voxels (LGNp) were defined as those in the P subdivision with strong activations in the chromatic localizer, while M-biased voxels (LGNm) were defined as those in the M subdivision with strong activations to the moving dots against black background in Exp. 2. The ROI-averaged BOLD timecourses in the contralateral LGNs clearly showed a transient response followed by a flat portion of sustained response (indicated by light and dark shaded areas in Fig. 2e and 2f, respectively). To avoid the non-specific transient signals due to changes in arousal and transient attention (Saalmann & Kastner, 2011), we focused on the sustained response period in the following analysis (see Fig. S2 for the results of the transient response). The time window of sustained response was defined from the ‘knee’ point on the timecourse to the stimulus offset. A leave-one-subject-out procedure was used to determine the sustained response period (see Methods).

The bar plots in Fig. 2e and 2f show the mean BOLD response in the sustained response period. Color-based attention exhibited distinct patterns of response modulation in the parvo- and magnocellular subdivisions of the LGN (significant three-way interaction among layer (LGNp/LGNm), attention (attended/unattended), and similarity (same/different) in a repeated measures ANOVA, F(1,24) = 5.934, p = 0.023, η_p_^2^ = 0.198). Follow-up tests showed a significant global effect of feature-based attention in LGNp (similarity × attention interaction: F(1,24) = 9.874, p = 0.004, η_p_^2^ = 0.291), but not in LGNm (F(1,24) = 0.001, p = 0.972, η_p_^2^ = 5.126×10^−5^). In LGNp, the BOLD response to stimuli presented at the unattended location was significantly enhanced when its color matched the attended color (“same” condition, t(24) = −2.663, p = 0.014, Cohen’s d = 0.565) compared to the “different” condition, while the BOLD response to stimuli presented at the attended location was significantly reduced (t(24) = −2.153, p = 0.042, Cohen’s d = 0.383). The statistics in LGNp was robust to the number of voxels when defining the ROI (Fig. S3). These findings clearly demonstrate that the similarity between the attended color and the color of unattended stimuli across the visual field significantly modulated BOLD responses in the P subdivision of the LGN. Our results cannot be explained by the difference in task difficulty between the same and different conditions, nor a “leakage” of spatial attention resource to the unattended side, since there was no significant difference in task performance (t(24) = 0.030, p = 0.976, BF_10_ = 0.211), and the response reduction of attended stimuli in the “same” condition was only observed in LGNp, but not in the cortex (Fig. 5d, see below).

The spatially global effect of color-based attention in the LGN was further demonstrated by a multi-voxel pattern analysis. We normalized each subject’s volumetric data in a standard space, and trained linear support vector machines (SVM) to predict whether a trial was from the “same” or “different” condition, using a leave-one-subject-out cross validation procedure (see Methods). The decoding accuracy for the LGN contralateral to the unattended stimuli was significantly above chance level (62%, p < 0.001, and p = 0.006 after correction for multiple comparison corrections across ROIs and attention conditions). Therefore, consistent with the univariate results, multivariate decoding shows that paying attention to a color matching the unattended stimuli across the visual field significantly modulated activity in the LGN contralateral to the unattended stimuli. Same vs. Diff pattern decoding was not significant in the SC and pulvinar, likely due to lower SNR in these regions.

### Color similarity modulation of attention is significant in the deeper layers of the SC

The SC is a laminated nucleus on the brainstem that plays importation roles in the controls of attention and eye movements. Neurons in the superficial layers of the SC are visuosensory, while those in the deeper (intermediate and deep) layers are visuomotor and motor. Since the superficial layers is about 1/3 of the thickness of the primate SC (May, 2006), we generated a normalized depth map in the SC and divided it into superficial (SCs) and deeper (SCd) ROIs (Fig. 3a, Fig. S4). Our results show that visual stimuli mainly activated SCs (Fig. 3b-d), which is consistent with the fact that neurons in the superficial layers mainly process visuosensory information. In Fig. 3c, there was a stronger activation in same compared to different condition in the SC contralateral to the unattended visual field (same - diff, CU).

**Figure 3.**
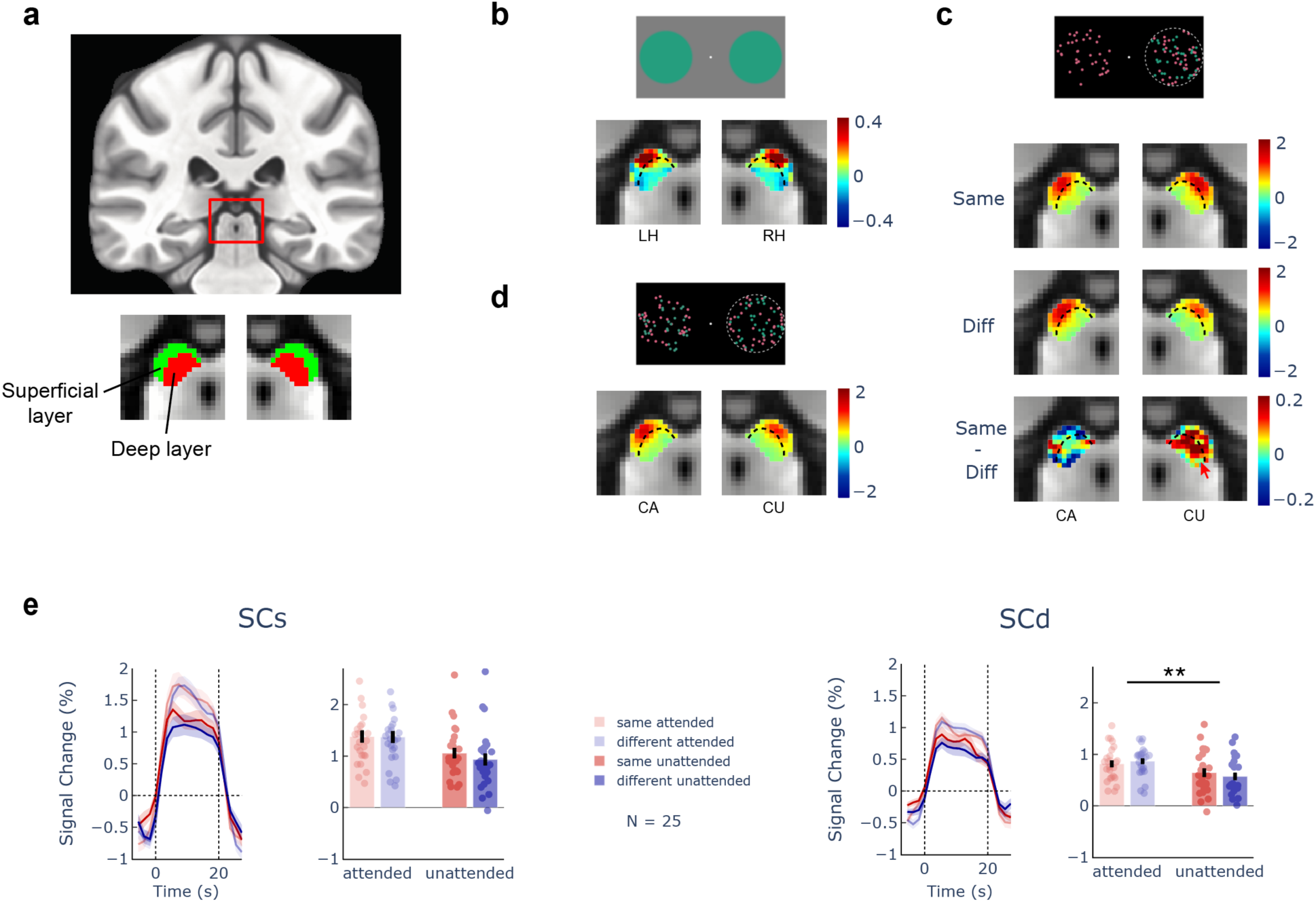
BOLD activation maps and ROI-averaged responses in the SC. (a) The location of SC in MNI space and the superficial and deeper layer ROIs. (b) (c) (d) are activation maps for chromatic localizer, Exp. 1, and Exp. 2, respectively. The red arrow in (c) highlights the activation to spatially global effect of color-based attention. Color bars indicate the BOLD response in percent signal change. LH, left hemisphere. RH, right hemisphere. CA, contralateral to the attended side. CU contralateral to the unattended side. (e) (f) are bar plots in superficial and deep layer respectively. Shaded areas and error bars indicate SEM across participants. **: p < 0.01.

The BOLD timecourse of the SC didn’t show a distinction in transient and sustained responses (Fig. 3e, 3f), so we averaged the response from 4s after the stimulus onset to the stimulus offset for statistical analysis. Only SCd showed a significant color similarity modulation of attention (attention × similarity interaction: F (1,24) = 7.753, p = 0.010, η_p_^2^ = 0.244). An insignificant trend was found in SCs (F (1,24) = 1.617, p = 0.216, η_p_^2^ = 0.063). Given the important role of the deeper layers of the SC in attention control, our results suggest that they are involved in the control of the spatially global effect of feature-based attention.

### Ventrolateral pulvinar shows a marginal effect of color similarity modulation of attention

The pulvinar is a large second-order thalamic nuclei that consists of several subregions with distinct functions and connectivity with cortical regions. According to a pulvinar atlas based on task co-activation patterns (Barron et al., 2015), we divided the posterior pulvinar into two ventral and two dorsal subregions (Fig. 4a, Fig. S5). The ventrolateral (vlPul) and ventromedial (vmPul) pulvinar primarily connect with the ventral and dorsal visual streams (Kaas & Lyon, 2007), respectively and selectively respond to parvocellular (P) and magnocellular (M) visual stimuli (C. Qian et al., 2023). The dorsal subregions are more involved in higher order cognitive functions and mainly connect with frontoparietal regions (Arcaro et al., 2018). Since our stimuli mainly activated the ventral pulvinar (Fig. 4b-d), we focused our analyses on the two ventral subregions. From the activation map in Fig. 4c, the vlPul contralateral to the unattended visual field shows stronger activations in the same compared to the different condition. Similar to the LGN, the timecourses of ventral pulvinar showed a clear transient response followed by a sustained response (Fig. 4e, 4f). The sustained response period showed a trend of color-similarity modulation of attention (similarity × attention interaction: F(1, 24) = 3.387, p = 0.078, η_p_^2^ = 0.124). A similar trend was also found for the transient response (Fig. S2).

**Figure 4.**
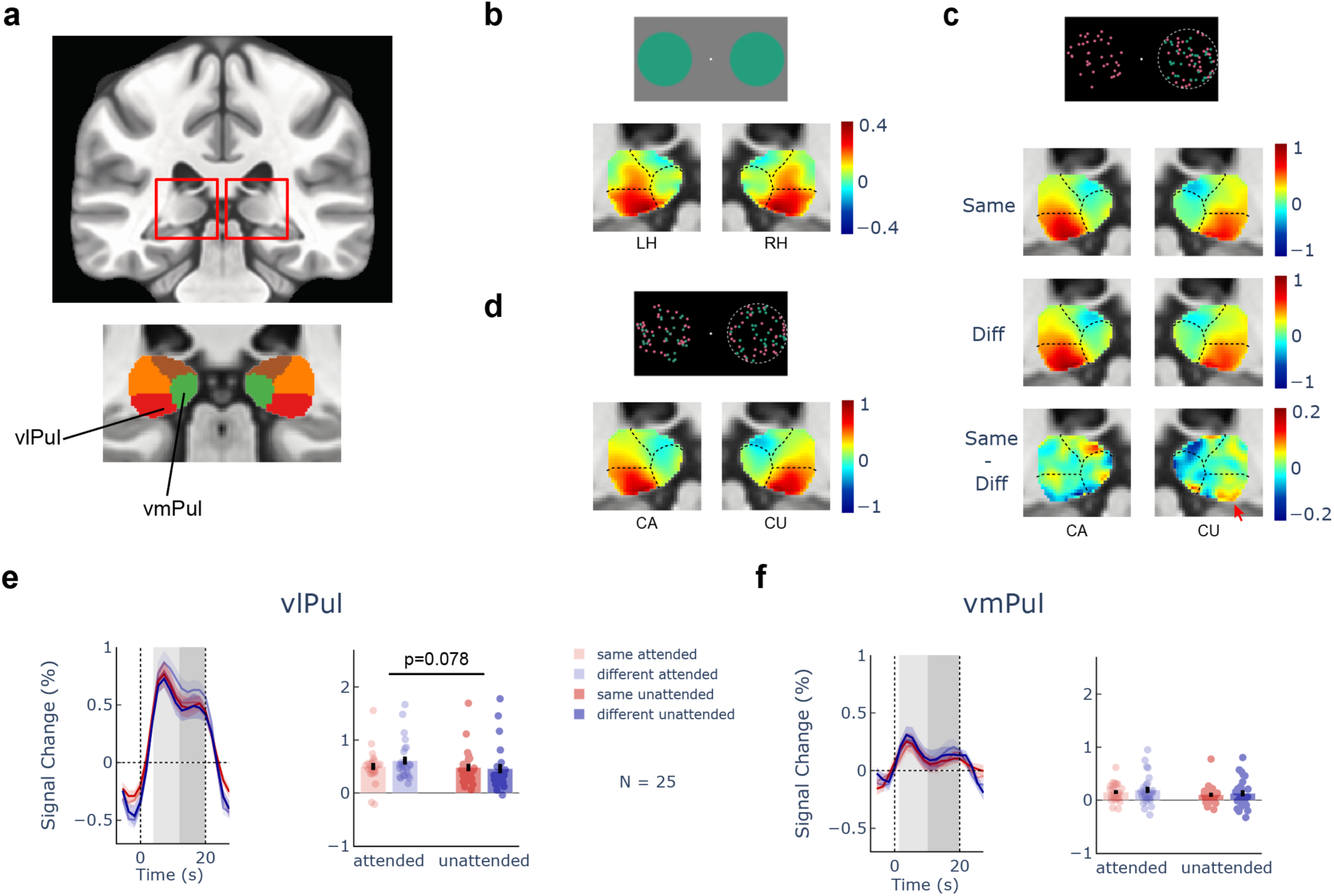
BOLD activation maps and ROI-averaged responses in the pulvinar. (a) The location of pulvinar in MNI space and its subdivisions. (b) (c) (d) are activation maps for the chromatic localizer, Exp. 1, and Exp. 2, respectively. The red arrow in (c) highlights the activation to spatially global effect of color-based attention. Color bars indicate the BOLD response. LH, left hemisphere. RH, right hemisphere. CA, contralateral to the attended side. CU contralateral to the unattended side. (e) (f) shows the ROI-averaged timecourses and bar plots in vlPul and vmPul. Light and dark shaded areas indicate the transient and sustained response period, respectively. Shaded areas and error bars represent SEM across participants.

### Spatially global effects of color-based attention in early visual cortices and frontoparietal regions

Fig. 5c shows the BOLD activation maps in cortical brain regions in Exp. 1. The early visual cortices and frontoparietal regions were activated in both the same (Fig. 5a) and different (Fig. 5b) conditions. The difference between the same and different conditions shows activations in V1, V4, and the posterior intraparietal sulcus (pIPS) in the hemisphere contralateral to the unattended stimuli. Cortical ROIs were defined by the functional localizer on the cortical surface (Fig. S6, S7). The ROI-averaged responses in Fig. 5d show significant global effects of color-based attention in the early visual cortices (V1-V4) and the pIPS. The timecourses for these cortical regions are shown in Fig. S8. Consistent with previous studies (Saenz et al., 2002; X. Zhang et al., 2018), we found significantly stronger global effect of color attention in hV4 than in hMT+ (similarity × ROI interaction in the unattended condition: F(1,24) = 23.858, p = 5.57×10^−5^, η_p_^2^ = 0.499). These results support the spatially global effects of feature-based attention in the early visual cortices and the posterior parietal cortex.

**Figure 5.**
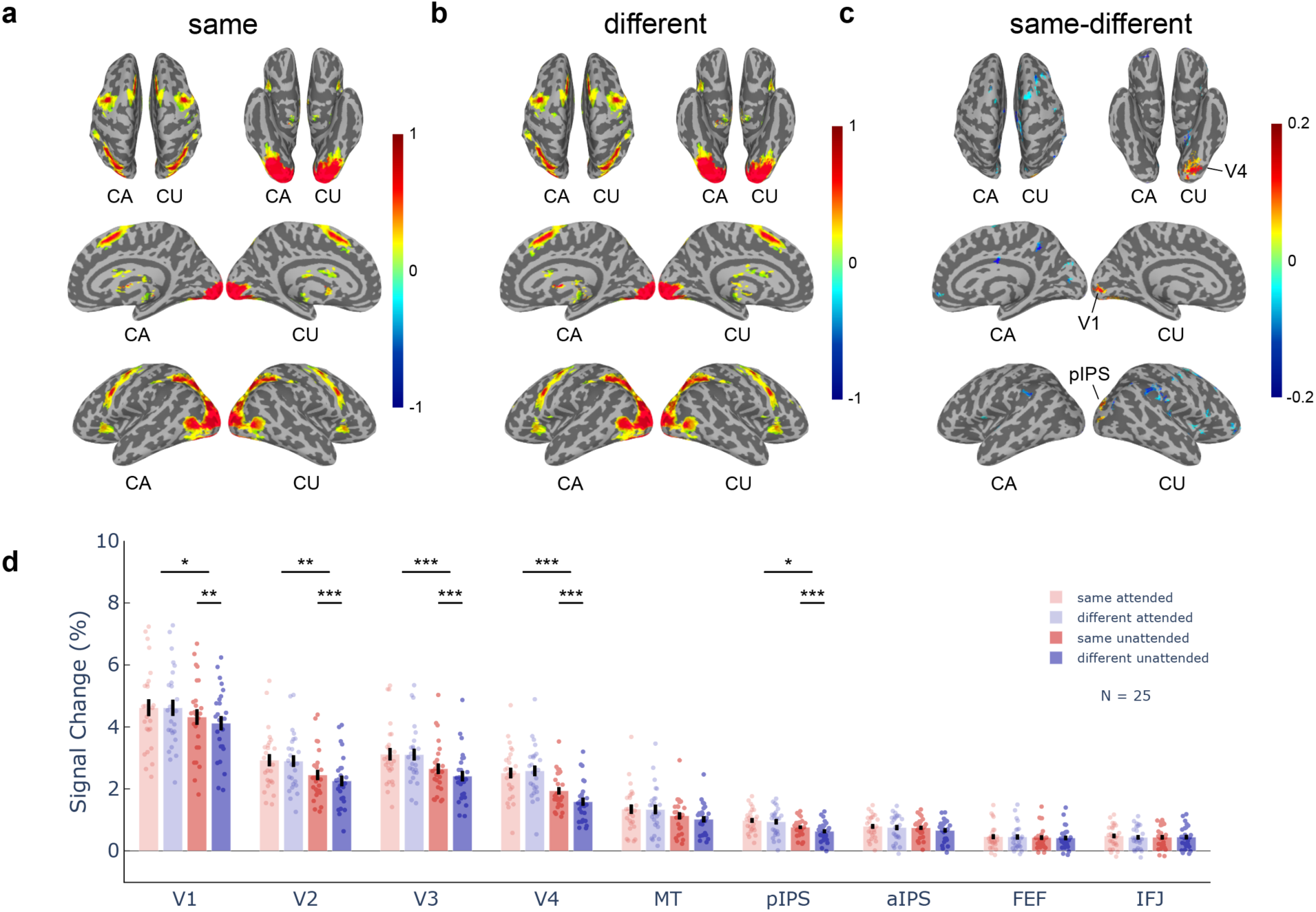
BOLD activation maps and spatially global effects of color-based attention in cortical regions. (a) (b) (c) are activation maps in same, different and same-different conditions respectively. *p* < 0.05, uncorrected. CA, contralateral to the attended side. CU, contralateral to the unattended side. (d) Bar plots in cortical ROIs. Error bars indicate SEM across participants. *: p < 0.05, **: p < 0.01, ***: p < 0.001, after Holm-Bonferroni correction. Abbreviations: pIPS, posterior IPS; aIPS, anterior IPS; FEF; frontal eye field; IFJ; inferior frontal junction. Error bars represent SEM across participants.

### Color-based attention modulates connectivity among cortical and subcortical regions

To further investigate how color-based attention modulates the information flow between cortical and subcortical regions, we used the dynamic causal modeling (DCM) to estimate the effective connectivity of fMRI signals among key brain regions as suggested by the previous analysis, including V1, V4, pIPS, LGN, SC and vlPul. A full DCM model was defined based on the empirical evidence of anatomical connections among these regions (Arcaro et al., 2015; Ghodrati et al., 2017; May, 2006; Shipp, 2003). In this model (Fig. 6a), the LGN receives the driving input; fixed or intrinsic connections were defined between and within brain regions; color similarity between the attended and unattended stimuli could modulate the effective connectivity in these connections, which were estimated separately in the attended and unattended conditions. At the first level, the full DCM model was estimated for the fMRI data within each hemisphere for each participant, and the results were averaged across the two hemispheres (Zeidman, Jafarian, Corbin, et al., 2019). The second level of analysis used Bayesian model reduction, Bayesian model average and parametric empirical Bayes to make inferences about the model evidence and connectivity strength (Zeidman, Jafarian, Seghier, et al., 2019).

**Figure 6.**
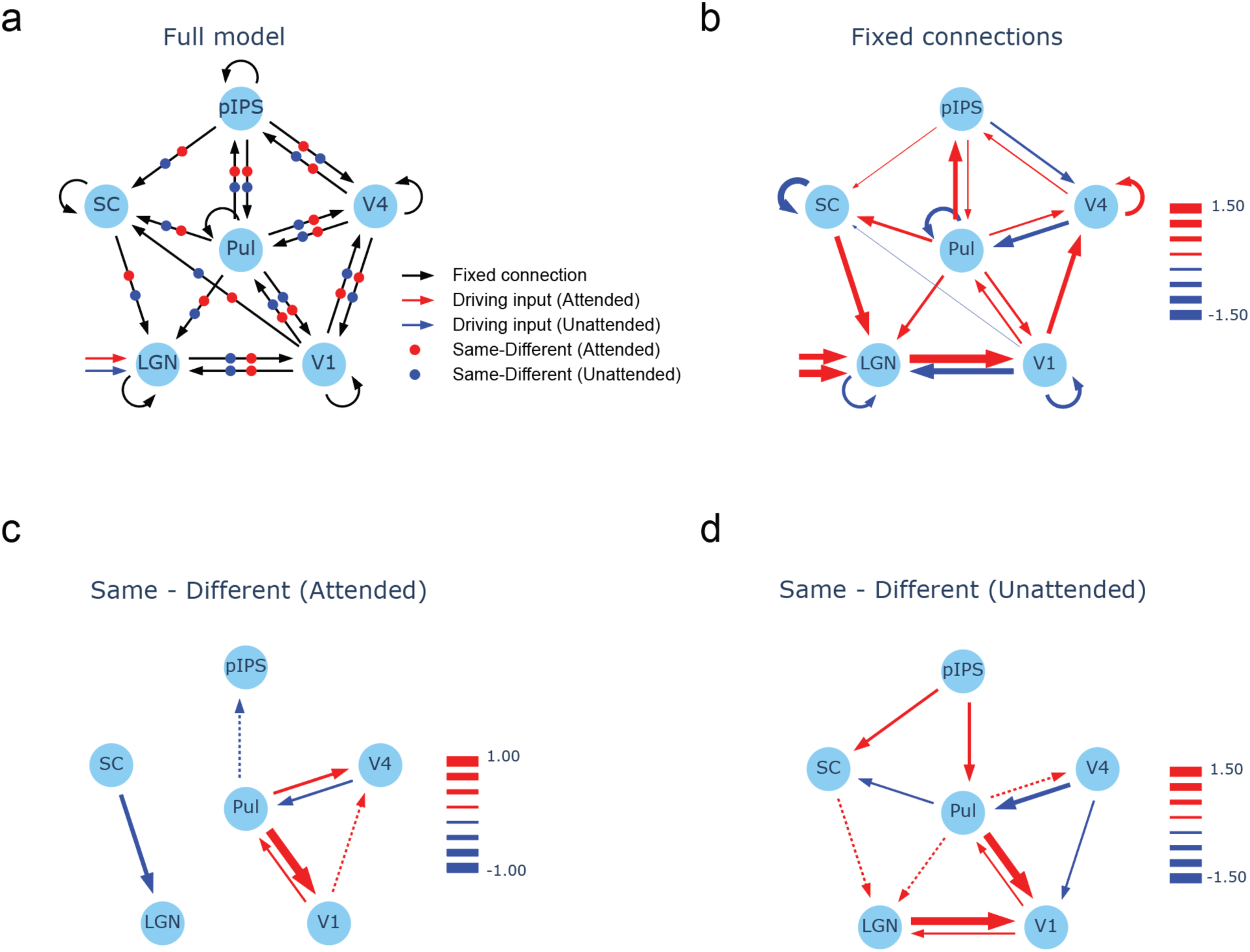
Dynamic causal modeling of effective connectivity in cortical and subcortical regions. (a) shows the full DCM model. (b) second-level results of fixed connections including the driving inputs. (c) (d) show the modulatory effect of feature similarity (same - diff) in attended and unattended conditions. Red and blue arrows in (b-d) denote positive and negative connections respectively. Dotted and solid lines represent connections above 0.75 and 0.95 posterior probabilities, respectively. The scale bar of line width indicates the connectivity strength (Hz).

Results of the second-level analysis are shown in Fig. 6b-d. Most of the fixed connections were highly significant (Fig. 6b), suggesting a good fit of the DCM model to our data. In the attended condition (Fig. 6c), color similarity (same vs. diff) with the unattended stimulus significantly modulated the connection from the SC to the LGN, and the connections between the pulvinar and early visual cortices. In the unattended condition (Fig. 6d), color similarity with the attended stimulus significantly enhanced the feedforward and feedback connections between the LGN and V1; LGN activity was also modulated by top-down signals from the SC and pulvinar originating from the pIPS; similar to the attended condition, the pulvinar also showed significant reciprocal modulatory connections with the early visual cortices. These results suggest that the spatially global effect of feature-based attention enhances both feedforward and feedback connectivity between the LGN and V1, modulated by top-down signals the parietal cortex through the SC and pulvinar.

### Attended color can be decoded from multi-voxel response patterns in the early visual cortices and frontoparietal regions

To further investigate whether the attended color can be decoded from brain activity to spatially overlapping red and green dots (from the attended location in Exp. 1 and Exp. 2, or from the unattended location in Exp. 2), we trained linear SVM classifiers with multi-voxel patterns of BOLD response in the hemisphere contralateral to the stimulus location. Results showed that in brain regions contralateral to the attended stimuli in Exp. 1 and Exp. 2 (Fig. 7 top panel, attended condition), the early visual cortices and frontoparietal regions showed significantly above chance decoding accuracy. In brain regions contralateral to the unattended stimuli in Exp. 2 (Fig. 7 bottom, unattended condition), V2, V4 and IPS also showed significantly above-chance decoding performance, suggesting spatially global effects of feature-based attention across the visual field. The lower decoding accuracy in the unattended condition is due to less training data compared to the attended condition (4 vs. 8 runs). Decoding results with 4 runs of data in Exp. 2 shows similar performance as those in the unattended condition (Fig. S9). For the subcortical regions, decoding performance was at chance level, likely due to their small size and low signal-to-noise ratio. These findings provide further support for the important role of frontoparietal regions in spatially global effects of feature-based attention.

**Figure 7.**
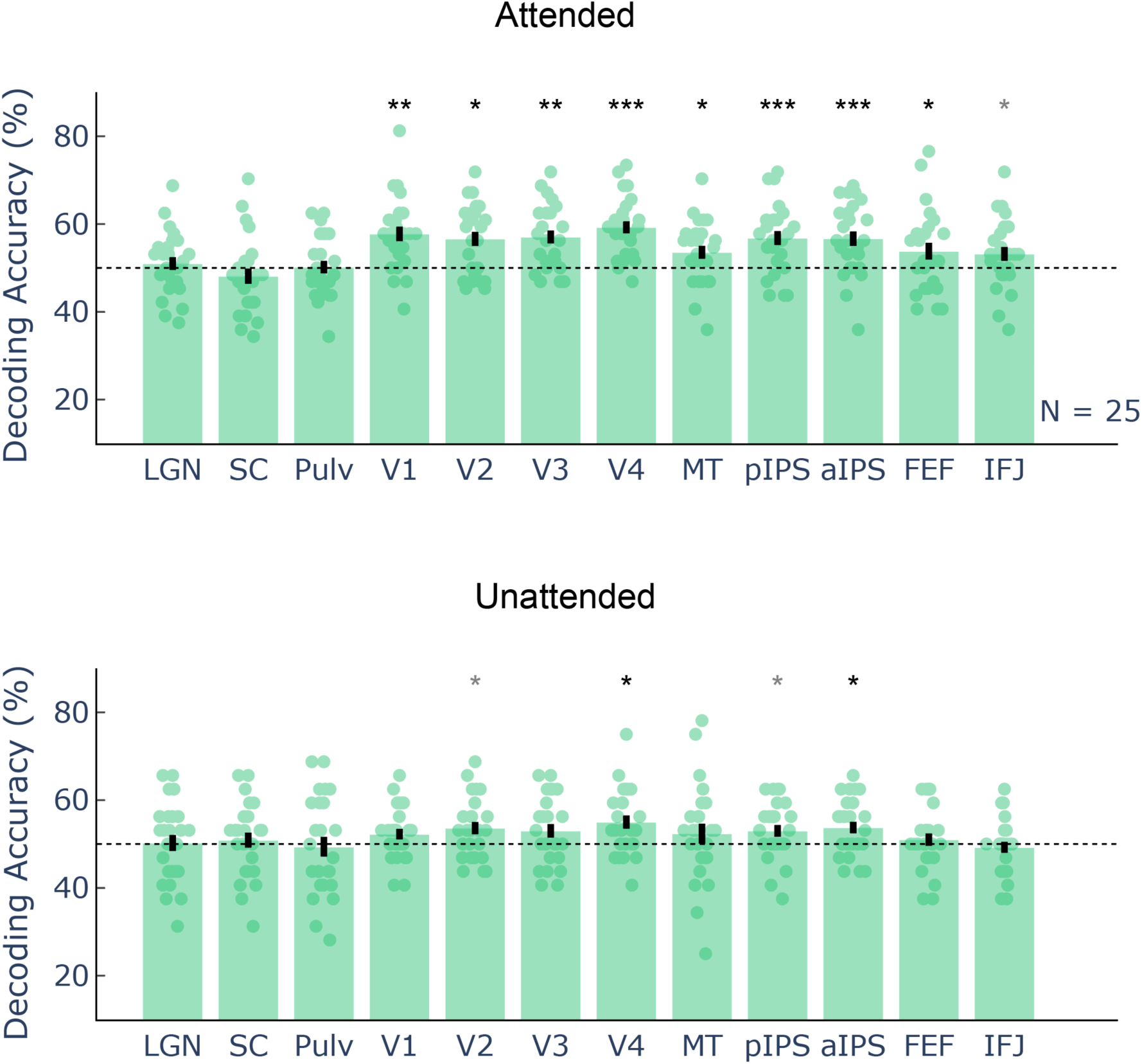
Decoding accuracy of attended color in brain regions contralateral to the attended and unattended locations. Top and bottom panels show the decoding accuracy of attended color in brain regions contralateral to the attended and unattended hemifields, respectively. Gray and Black asterisks indicate significance before and after correction for family-wise errors, respectively. *: p < 0.05, **: p < 0.01, ***: p<0.001. Error bars represent SEM across participants.

## Discussion

Using high-resolution 7T fMRI, we found spatially global effects of feature-based attention in functional subdivisions of human subcortical nuclei. When an unattended stimulus in the opposite visual field matched the color of the attended stimulus, BOLD signals in the P subdivision of the LGN were significantly enhanced for the unattended stimulus while reduced for the attended stimulus. The color similarity modulation of attention across the visual field was also found in the deeper layers of the SC, and showed a trend in the ventrolateral pulvinar. Effective connectivity analyses showed that the spatially global effect of color-based attention enhanced both feedforward and feedback connectivity between the LGN and V1, and was controlled by top-down signals from the IPS of parietal cortex through the SC and pulvinar. Attended color can be decoded from multi-voxel response patterns in frontoparietal regions in both hemispheres.

In an electroencephalographic (EEG) study, the spatially global effect of feature-based attention was found to modulate the early P1 component of event-related potentials in the occipital lobe around 100 ms (W. Zhang & Luck, 2009). Our high-resolution fMRI results demonstrate that feature-based attention enhanced both feedforward and feedback processing of the attended feature information throughout the visual field in the LGN of the thalamus (Fig. 2c, 2e; Fig.6d), which is the earliest stage of feature-selective processing after the retina. Furthermore, color similarity modulation was significant in the P but not in the M layers of the LGN, consistent with the feature similarity gain model of attention (Treue & Martinez Trujillo, 1999).

One interesting finding of the current study is that LGN activity to the attended stimulus was significantly reduced when the color of the unattended stimulus from the opposite visual field matched the attended color (Fig. 2e). A similar trend was also observed in the vlPul (Fig. 4e). One possible explanation is that the spatially global effect of feature-based attention comes at a cost of the attended stimulus processing. For example, when a stimulus at an unattended location shared the attended feature, the subcortical mechanisms might automatically spread attention to and sample information from that location, resulting in a slight reduction in response gain to the attended stimulus. This should reflect a change in the scope of attention or a switch to the global sampling strategy, rather than a distraction of spatial attention away from the attended location, since there was no significant difference in task performance between the same and different conditions. Another possibility is that a nearby stimulus sharing the attended feature produced a stronger suppression effect on the ignored stimulus feature at the attended location. This could lead to an overall reduction of BOLD signals in the thalamic nuclei contralateral to the attended location. To distinguish these two possibilities, it requires to isolate the processing of attended and unattended stimulus features at the attended location, which is not possible in the current study because the color decoding performance in subcortical regions was at chance level (Fig. 7). In contrast to the subcortex, cortical regions did not show a significant difference between the same and different conditions at the attended location (Fig. 5; Fig. S8), suggesting that the suppression effect on the response of the attended stimulus should reflect a subcortical mechanism. Also, this is unlikely due to difference in eye movement in the same and different conditions, since this would have influenced both cortical and subcortical regions.

Another key finding of our study is that color-based attention significantly modulated BOLD signals in the deeper layers (Fig. 3f), but not in the superficial layers (Fig. 3e) of the SC. The superficial layers of the primate SC mainly process visuosensory information, while the intermediate and deep layers controls attention and eye movement (May, 2006). In a single-unit study in monkeys (B. J. White et al., 2009), neurons in the intermediate SC can respond strongly to isoluminant color stimuli, with a longer latency compared to luminance response, suggesting a cortical influence from frontoparietal attention networks (Leichnetz & Goldberg, 1988; May, 2006). Thus, the SC may play a role in controlling the spatially global effect of feature-based attention. In support of with this claim, feature similarity between the attended and unattended stimuli across the visual field modulated the effective connectivity from the SC to the LGN (Fig. 6c, 6d), and the SC activity was also modulated by top-down connection from the IPS of parietal cortex (Fig. 6d). Although the spatially global effect of color-based attention was not significant in the pulvinar (Fig. 4), effective connectivity results suggest that it might play a role in regulating information transmission between cortical regions for information processing in both attended and unattended visual field (Fig. 6). Thus, the pulvino-cortical connections could be an important mechanism supporting the spatially global effect of feature-based attention.

In cortical regions, our results replicated the previous findings of spatially global effects of feature-based attention in the early- and intermediate-level visual cortices (Fig. 5) (Martinez-Trujillo & Treue, 2004; Saenz et al., 2002; Serences & Boynton, 2007; W. Zhang & Luck, 2009; X. Zhang et al., 2018). More importantly, we also found this effect in the posterior IPS of parietal cortex. Effective connectivity analyses showed that top-down signals from the IPS significantly modulated activity in lower-level brain regions through the pulvinar and SC in feature-based attention across the visual field (Fig. 6d). Furthermore, attended color can be decoded from multivoxel response patterns in the contralateral IPS in both attended and unattended conditions (Fig. 7). These findings support an important role of the IPS in top-down control of the spatially global effect of feature-based attention. In the frontal regions (e.g. FEF, IFJ), attended color can be decoded from contralateral hemisphere in the attended condition, consistent with a previous fMRI study at 3T (Liu et al., 2011). However, while another 3T fMRI study showed enhanced frontal activity in the same compared to different conditions (X. Zhang et al., 2018), we did not find this effect in our data (Fig. 5). One possible reason for this discrepancy is the difference in stimulus eccentricity. Due to the limited field of view of the head coil at 7 T, the eccentricity of our stimuli (4.25 degrees) is much smaller compared to the previous 3T study (8.5 degrees). Given the large neuronal receptive field in frontal cortex, these regions might be more involved in controlling global feature-based attention in a larger field of view. Another possible reason is that, the spatial attention effect, i.e., the response difference between the attended and ignored sides, was only about half of the effect size in (X. Zhang et al., 2018). This may result in a larger baseline activity on the ignored side and a ceiling effect, making the modulation by feature-based attention between the same and different conditions less evident.

To summarize, we found that feature-based attention modulated the processing of the attended feature throughout the visual field as early as in the LGN of the thalamus. Top-down signals from the parietal cortex play roles in controlling the effect of feature-based attention in the early visual areas through the SC and ventral pulvinar.

## Methods and Materials

### Participants

A total of 25 healthy human adults (15 females; 19-31 years of age) participated this study. All participants had normal or corrected-to-normal vision and reported no history of neuropsychological or vision disorders. The spatially global effects of feature-based attention showed a large effect size (d-prime > 0.8) in cortical regions (Saenz et al., 2002; X. Zhang et al., 2018). Due to the lower SNR in subcortical regions, the effect size was expected to be lower, thus we selected a sample size of 25 participants to provide 80% power to detect at least a medium effect size (d-prime = 0.5). The experimental procedures were approved by the ethical review board of Institute of Biophysics, Chinese Academy of Sciences. Written informed consent was obtained from all participants prior to their participation in the study.

### Stimuli and procedures

#### Apparatus and calibrations

Visual stimuli were generated in MATLAB (Mathworks Inc.) with Psychtoolbox Version 3 (PTB-3) (Kleiner 2007; Brainard 1997). In the behavioral session, visual stimuli were presented by a LCD monitor at 1920 × 1080 pixels and 120Hz refresh rate (Display++, Cambridge Research System). In the fMRI session, stimuli were presented by an MRI compatible LCD monitor at 1920 × 1080 pixels and 60Hz refresh rate (BOLDscreen 3D, Cambridge Research System). Participants viewed the stimuli through a rear-view mirror mounted inside the head coil. Both displays were gamma corrected to achieve a linear luminance output with lookup tables. The red, green, and blue spectral power distributions were measured using a Photo Research PR-655 spectroradiometer (Chatsworth, CA, USA).

#### Behavioral experiment

Spatially overlapping red and green dot fields were presented in a circular region on each side of the fixation point at 4.5° eccentricity (Fig. 1a). The size of the dot fields was 5.5° in diameter. Dots were updated at random locations within the circular aperture every 33 ms. The chromaticity of the stimuli was defined in a three-dimensional cone contrast space (Cole et al., 1993; Mullen et al., 2007). In this space, cone contrasts represent the ratios of cone excitation levels within the same cone type. Therefore, using cone contrasts allows for control of visual inputs from the early stage of color processing, namely, the absorption of light by cones as a function of wavelength. We used a red-green stimulus that isolated L-/M-cone opponency by setting the activation weights of each cone to 1, -a, and 0, where a was the ratio of L-to M-cone weights for red-green (RG) isoluminance. To determine RG isoluminance, participants performed a minimum motion task (Anstis & Cavanagh, 1983) before the experiments by adjusting a counter-phase flickering horizontal grating until a minimum motion was perceived.

Participants performed two independent color discrimination tasks simultaneously on both visual fields. The magnitude of the red/green color contrast change was determined for each participant in a pilot experiment using a 3-down-1-up staircase procedure. In the beginning of each trial in the formal experiment, two colored dots were presented on each side of the fixation point to indicate the target dots to be attended in the following presentation. After pressing a button, spatially overlapping red/green dots were presented twice on both sides of the fixation point in two 1000-ms intervals, separated by 100-ms black background. Participants’ task was to respond in which interval the target dots had higher red/green cone contrast. After responding to stimuli in both visual fields, feedback was given by a cross (wrong) or a checkmark (correct). The experiment consisted of four runs of cue combinations: red or green cues on both sides (same condition), left red/right green or left green/right red (different condition). 50 trials were collected for each run, and a total of 100 trials were collected for the same or different conditions. The order of the four runs was randomized across participants.

#### fMRI localizer

The localizer stimuli were uniform red/green disks alternating at 2Hz on each side of the fixation point against an equiluminant gray background, presented at the same size and location as those in the behavioral and fMRI experiments. The chromatic flickers were presented for 16 seconds in each stimulus block, followed by a 16-s fixation period. Each run lasted 272 seconds. Participants performed a simple fixation task, detecting occasional contractions of the fixation point. A total of 2 runs were scanned for each participant.

#### fMRI experiment 1

Red and green dot fields were identical to those used in the behavioral experiment. A black background was used to strongly activate both M and P layers of the LGN. Before the start of each run, participants were informed of the location of the stimulus to be attended by an arrow at the fixation point. Stimuli were presented in 20-s blocks, alternating with 12-s fixation blocks. From 2 s before the start of each stimulus block, the white fixation point turned red or green to indicate the target color to be attended to in the following presentation. Each stimulus block consisted of 8 trials. Spatially overlapping red and green dots were presented at the cued location, with a set of dots of either the same or different color as the attended dots were presented on the opposite side of fixation. The dots stimuli were presented twice in each trial, in two 833-ms intervals, separated by 83-ms fixation between the two presentations, followed by another 750-ms fixation before the next trial. Participants were required to maintain fixation and respond to which interval the attended dots had higher red/green cone contrast. Each run lasted 284 seconds. A total of 4 runs were collected for Exp. 1. The difficulty of the color discrimination task was adjusted to the threshold level by an adaptive staircase procedure during the structural scan.

#### fMRI experiment 2

Stimuli and procedures were identical as those in Exp. 1, except that spatially overlapping red and green dots were presented on both the attended and unattended sides of the fixation point. 4 runs of data were also collected for Exp. 2.

### MRI data acquisition

MRI data were collected on a 7T scanner (Siemens Magnetom) with a 32-channel receive 1-channel transmit head coil (NOVA Medical). Functional data were collected with a T2*-weighted 2D GE-EPI sequence (1.5 mm isotropic voxels with 68 axial slices, FOV = 183×183 mm, TR = 2000 ms, TE = 22 ms, flip angle = 70°, phase partial Fourier = 6/8, GRAPPA = 2, multiband factor = 2, phase encoding direction from A to P). T1-weighted anatomical volumes were acquired with a MP2RAGE sequence (0.7 mm isotropic voxels, 256 sagittal slices, FOV = 224×224 mm, GRAPPA = 3, TR = 4000 ms, TE = 3.05 ms, TI1 = 750 ms, flip angle = 4°, TI2 = 2500 ms, flip angle = 5°, phase/slice partial Fourier = 7/8). A total of 10 runs of fMRI data were scanned in the 7T experiment, including 2 runs for the chromatic localizer, and 4 runs each for Exp. 1 and Exp. 2 (one participant finished 3 runs for Exp. 2 due to physical discomfort). The order of experimental conditions was counterbalanced across runs and participants. A bitebar was used to reduce head motion.

### MRI data preprocessing

T1-weighted MP2RAGE anatomical images were segmented and reconstructed into gray matter and white matter surfaces in FreeSurfer (version 6.0) (Fischl, 2012). GE-EPI functional images were preprocessed by AFNI (Cox, 1996), advanced normalization tools (ANTs) (Avants et al., 2011), and custom Python code, in the following steps: slice timing correction, EPI distortion correction with nonlinear warping (Blip up/down method), rigid body motion correction, alignment of corrected EPI images to T1-weighted anatomical volume (cost function: lpc), upsampling by a factor of 2 to reduce the resolution loss caused by interpolation (Wang et al., 2022; P. Zhang et al., 2015), and per run scaling as percent signal change. To minimize image blur, all spatial transformations and upsampling were combined and applied to the functional images in one interpolation step (sinc method). General linear models with a short time-to-peak HRF (Block4 in AFNI) were used to estimate BOLD signal change from baseline for each stimulus condition. Motion parameters were included as regressors of no interest. The statistical results of subcortical regions were transformed into MNI space using a nonlinear mapping (antsRegistration) between the anatomical volume and the symmetric T1-weighted template from CIT168 atlas (Pauli et al., 2018). To improve the registration accuracy in subcortical regions, a spherical weight mask including the visual thalamus and the upper brainstem was used in the nonlinear warping process.

### ROI definition and analysis

Anatomical ROI for subcortical regions were first defined on the high-resolution MNI template, and then manually edited for each individual based on their T1-weighted anatomical images in MNI space. With high T1 contrast at 7 T (Fig. S10), the LGN appeared darker than the surrounding white matter and brighter than the cerebrospinal fluid in the ventricle, allowing its anatomical boundaries to be accurately defined by an experienced experimenter. Due to their proximity, the LGN and the ventral pulvinar of the thalamus were carefully delineated on the sagittal slices. To assist the group-level analysis, the coordinates of the left LGNs were mirror-flipped to the right, and then all LGNs were registered to the LGN of the MNI template by a twelve-parameters linear (affine) transformation. SC and pulvinar ROIs were defined using the same procedure.

To analyze data from the functional subdivisions of the LGN, a normalized layer index was calculated for each voxel in the MNI template (Fig. S1). Two layers of voxels were first defined corresponding to the ventral and dorsal surfaces of the LGN. For the rest of voxels, we calculated a layer index as the normalized distances to the dorsal and ventral surfaces (depth = 0 and 1 correspond to the dorsal and ventral surfaces, respectively). ROIs for the M and P subdivisions were determined from the layer index map according to the volume ratio of M/P layers of human LGNs (M/P volume ratio = 1/4) (Andrews et al., 1997). To generate the depth map of the SC (Fig. S4), two layers of voxels were first defined from the superficial and deep surface of the SC. Then a normalized depth map was calculated for each voxel as the ratio of shortest distances to the superficial and deep surfaces (depth = 0 and 1 correspond to superficial and deep surfaces, respectively). Based on the anatomy of the primate SC (May, 2006), the SC was split into a superficial and a deeper layer compartment at depth = 1/3. Pulvinar subdivisions were delineated based on task-coactivation patterns (Fig. S5) (Barron et al., 2015), including the ventromedial (vmPul), ventrolateral (vlPul), dorsolateral (dlPul), dorsomedial (dmPul) and anterior (aPul) subdivisions of pulvinar. Since our stimuli mainly activated the ventral pulvinar (Fig. 4c), only data from vlPul and vmPul were used for the ROI-based analysis.

Cortical ROIs were defined on the inflated standard surface (std141 in AFNI). The early visual cortices V1, V2, V3, V4 and MT were defined based on the polar angle atlas from the 7T retinotopic dataset of Human Connectome Project (Fig. S6) (Benson et al., 2014, 2018). Frontoparietal areas pIPS, aIPS, FEF and IFJ were defined manually based on the anatomical landmarks and the group-level activations in the functional localizer (thresholded at t > 0, Fig. S7). Volume ROIs were generated by transforming the surface ROIs to the volume space (3dSurf2Vol in AFNI).

For the analysis of ROI-averaged responses and timecourses in Exp. 1, 24 upsampled voxels (corresponding to the volume of 3 original voxels) that are most responsive to the chromatic localizer were selected from the P subdivision of the LGN, while the same number of voxels were selected from the M subdivision with the strongest activations to the attended dots on black background in Exp. 2. For the superficial and deep SC, voxels (n = 24) most responsive to the attended stimulus in Exp. 2 were selected from the ROIs. From vmPul and vlPul subdivisions, 100 voxels that are most responsive to the attended stimulus in Exp. 2 were selected. For the early visual cortices, voxels with significant activation to the chromatic localizer (p < 0.05 uncorrected) were used. All voxels were used in the frontoparietal ROIs.

The timecourses of LGN and pulvinar showed a clear transient response followed by a sustained phase (Fig. 2 and Fig. 4). to avoid the non-specific transient signals due to arousal change and transient attention (Saalmann & Kastner, 2011), we focused on the sustained response in the analysis of ROI-averaged responses. The time window for the sustained response was defined from the “knee” point, marking the transition between the transient and sustained phases up to the stimulus offset. A leave-one-subject-out approach was employed to determine the sustained response time window based on the group-averaged timecourse. Different stimulus and attention conditions shared the same baseline for the timecourse analysis, which was estimated by a general linear model.

### Attended color decoding

For both cortical and subcortical areas, we also trained linear support vector machine (SVM) classifiers (Pedregosa et al., 2011) in the native space to decode the attended color (red or green) from multivoxel response patterns to stimulus blocks in ROIs contralateral or ipsilateral to the attended location, using a leave-one-run-out cross-validation procedure. Each stimulus block was modeled with a separate regressor in the GLM. For attended color decoding in cortical regions, feature selection was performed with K-1 runs of training data to select voxels with strong color selective responses in the ROI (top 10% activated voxels in the chromatic localizer). Activation patterns were normalized by z-score across samples (or stimulus blocks). Given the limited number of voxels in subcortical ROIs, all voxels were used in this analysis. An SVM classifier was trained from K-1 runs of data, and the distance to the decision boundary was used to classify the attended color for each stimulus block from the remaining run. The cross-validation procedure was repeated for all runs and the results were pooled together to calculate the decoding accuracy.

### Color similarity (same vs. different) decoding

Since subcortical regions exhibit less variability in anatomy, we normalized the volumetric data of each subject in the MNI space, and trained linear support vector machines to predict whether a trial was from the “same” or “different” condition, using a leave-one-subject-out cross validation (i.e., we trained on N-1 subjects, and tested on the unseen one, assuming that the difference in activity patterns between the two conditions was consistent across subjects). The analysis was performed separately for the subcortical ROIs contralateral to the attended or unattended side. Feature selection was done by choosing voxels with largest activation difference between the two conditions, and the fraction was determined using a nested 4-fold cross validation within the training set. Each data sample was then preprocessed by dividing its L2-norm to remove the effect of overall activation level difference. The test samples were first averaged by category to boost SNR before being fed to the classifier. The critical level for evaluating the significance of decoding accuracy corrected for multiple comparisons was obtained in a permutation procedure. The null distribution for the maximum mean decoding accuracy was constructed by randomly flipping the sign of accuracy difference away from the chance level for each subject, taking the group mean, and finding the maximal value across conditions and ROIs. This was repeated for 1000 times and the accuracy at 5% was the critical value.

### Dynamic causal modeling

Effective connectivity of fMRI data was analyzed using the DCM module of SPM12 (version 2020-Jan-13th). fMRI data were preprocessed in AFNI with slice timing and motion corrections. The mean timecourses of the LGN and SC were averaged from 24 upsampled voxels with the strongest similarity-by-attention interaction. The vlPul timecourses were averaged from 100 voxels with the strongest interaction effect. ROIs for the cortical regions were defined as voxels with significant feature similarity effect (same-different t > 1.96) in the unattended condition. A full DCM model was defined based on the empirical evidence of anatomical connections among these brain regions (Fig. 6a). At the first level, the full DCM model was estimated for the fMRI data within each hemisphere for each individual, and the results were averaged between the two hemispheres using Bayesian fixed effect (FFX) average (spm_dcm_average). At the second level, we used Bayesian model reduction, Bayesian model average and parametric empirical Bayes to make inference about the model evidence and connectivity strength (spm_dcm_peb and spm_dcm_peb_bmc).

### Statistics

For the ROI-averaged responses, we performed repeated measures (rm) ANOVA with attention (attended/unattended), feature similarity (same/diff) and ROI as the fixed effects, and subjects as the random effect. Post-hoc paired t-tests were performed if there was a significant attention by similarity interaction. Based on the rationale of the seminal paper by Levin and colleagues (Levin et al., 1994), test of simple effects following a significant 2-by-2 interaction does not need further correction for family-wise errors. Holm-Bonferroni correction was used to control the family-wise error of the attention by similarity interaction effect across ROIs in Fig. 5d. A non-parametric permutation test was used to control the family-wise error of attended color decoding performance across ROIs (Fig. 7) (Nichols & Holmes, 2001). In each permutation, the label of attended color was randomly shuffled within each run, and the entire cross-validation was applied, resulting a group mean accuracy for each ROI under null hypothesis. The permutation procedure was repeated 1000 times. A null distribution was generated by taking the maximum group-level accuracy across ROIs in each permutation. The family-wise error of the decoding performance in each ROI was derived from the null distribution.

## Data and code availability

Data and code to reproduce the main findings of this study will be made available upon publication.

## Acknowledgements

This study was supported by STI2030-Major Projects (2022ZD0211900, 2021ZD0204200, 2022ZD0204802), National Natural Science Foundation of China (31930053), and Chinese Academy of Sciences (2021089).

## Supplementary figures

**Figure S1.**
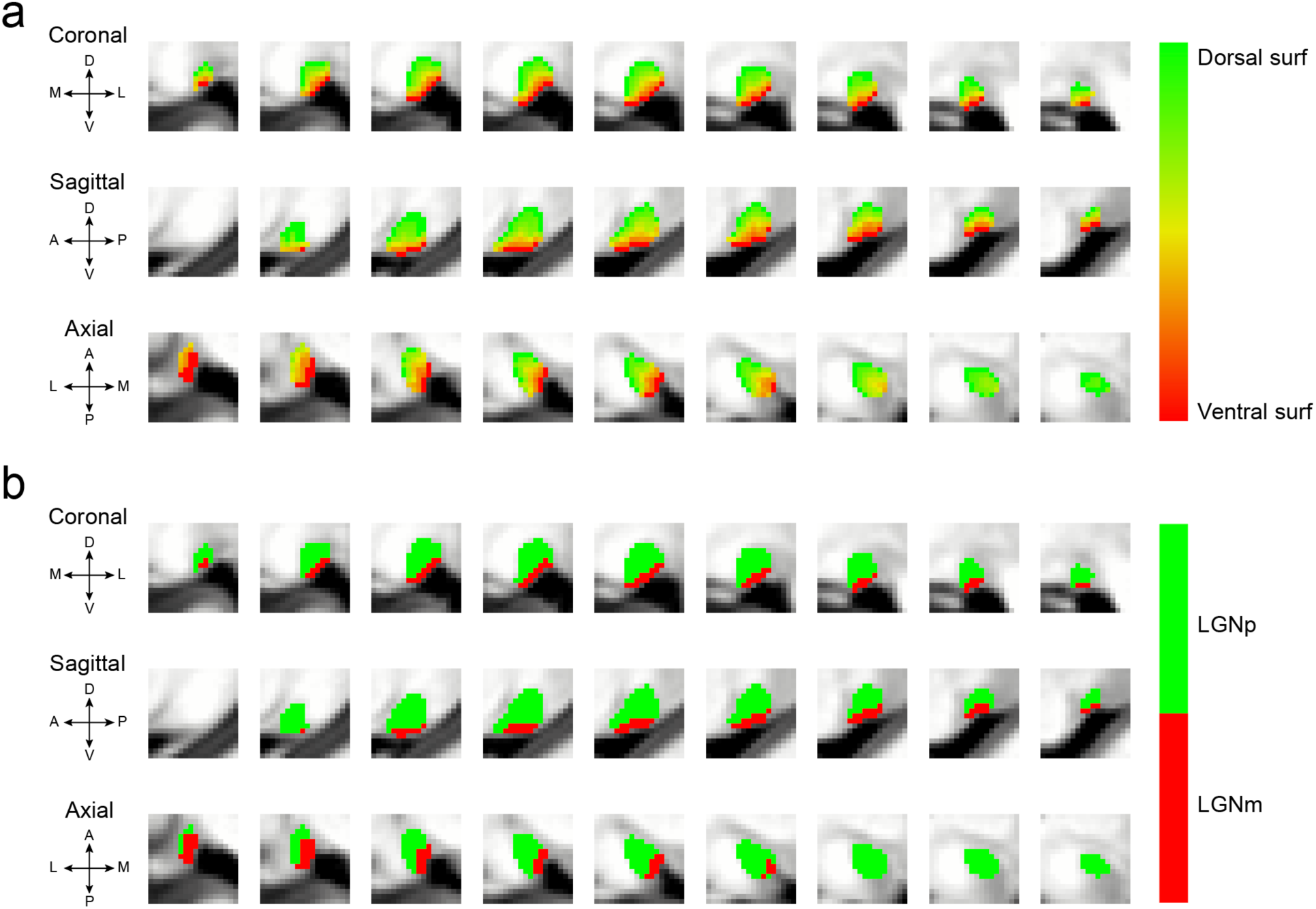
Normalized layer index map and LGN subdivisions. LGNp: parvocellular subdivision of the LGN, LGNm: magnocellular subdivision of the LGN.

**Figure S2.**
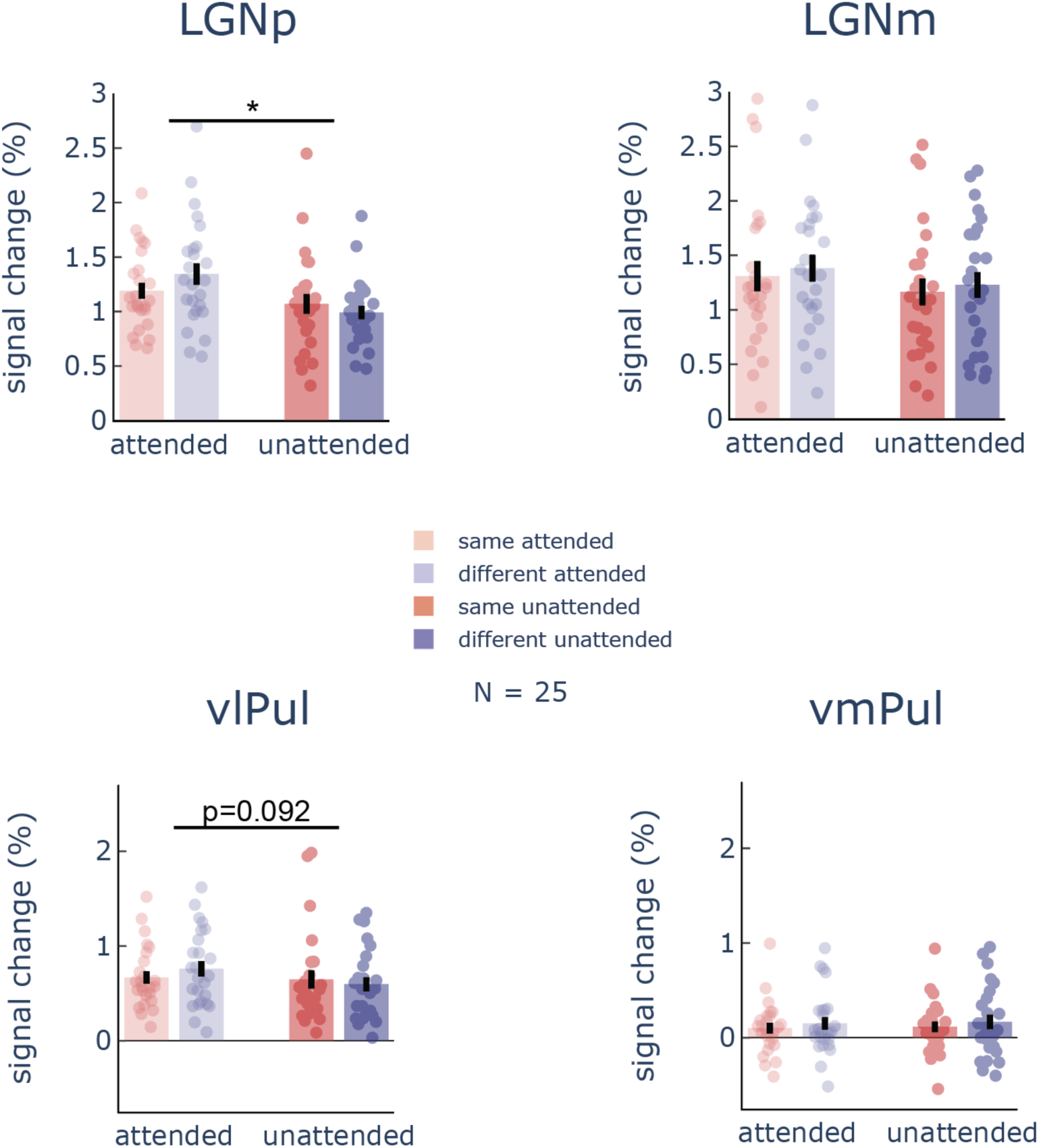
ROI-averaged responses of the transient period in the LGN and ventral pulvinar. The Asterisk and p value above the horizontal black lines indicate statistical signficance of attention (attended/unattneded) by similarity (same/diff) interaction. Errorbars represent ±SEM.

**Figure S3.**
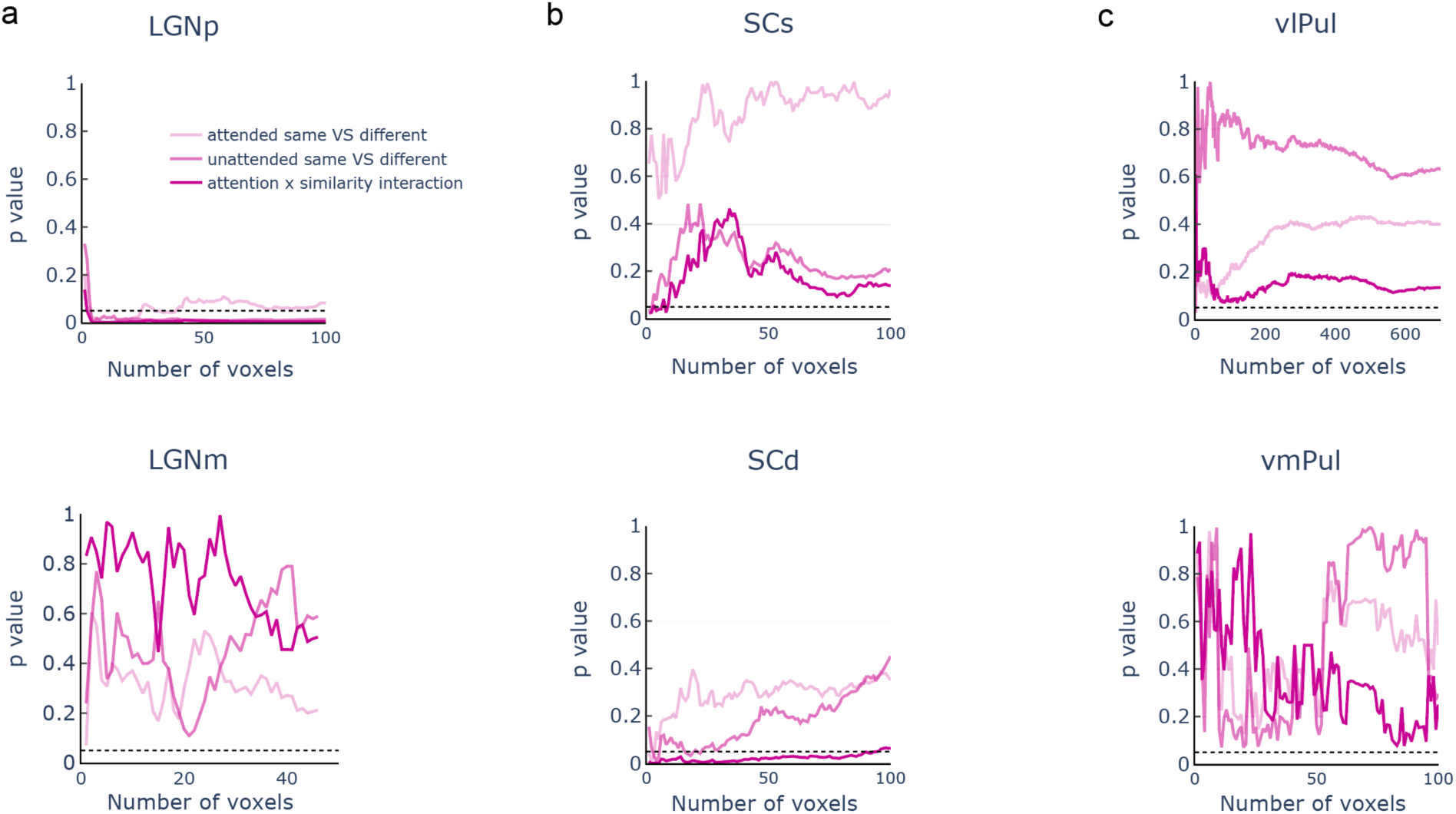
Statistical significance as a function of the number of voxels in the subcortical ROIs. The line plots show the p values attention by similar interaction, and post-hoc ttest of the difference between same and different in the attended and unattended conditions.

**Figure S4.**
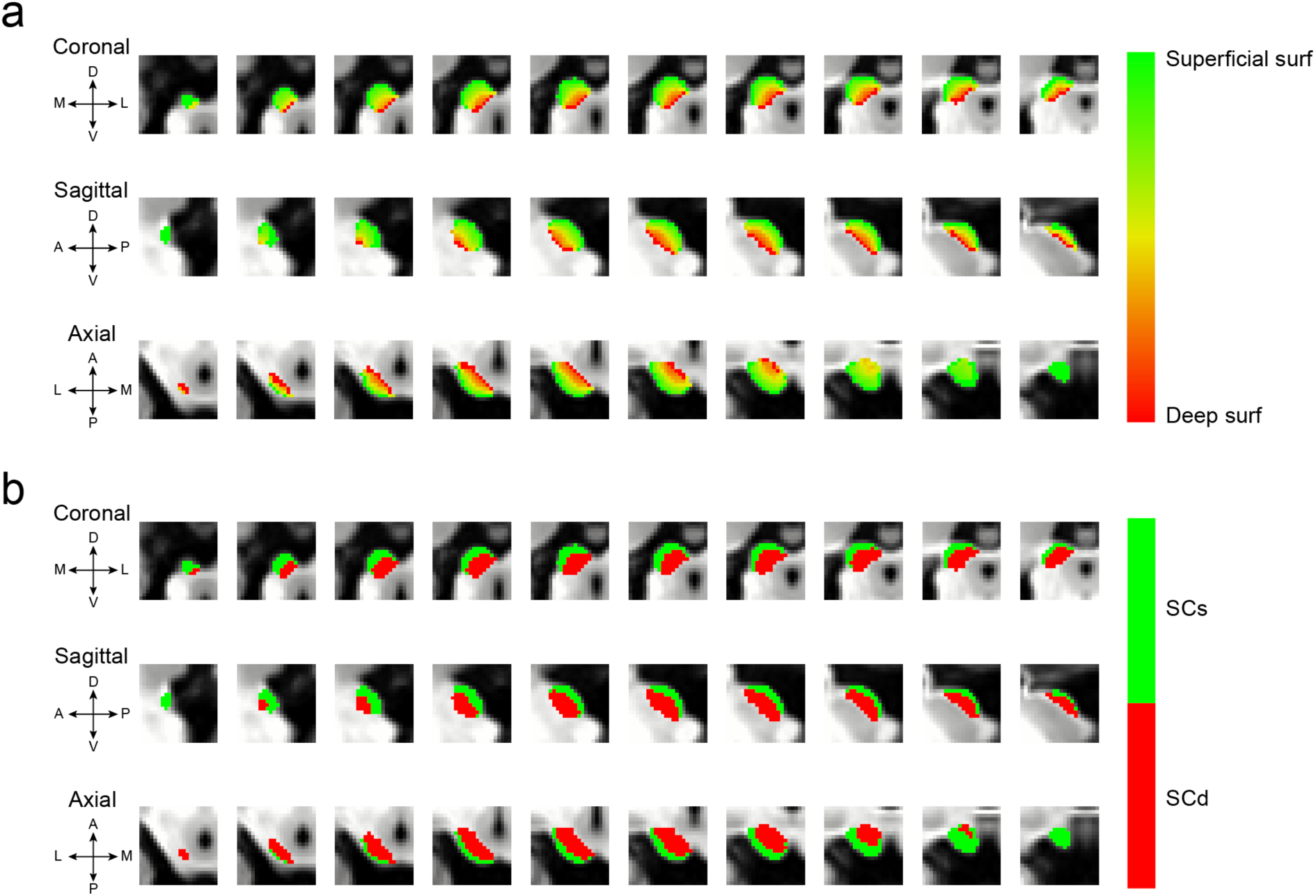
Normalized layer index map and subdivisions of the SC. SCs: superficial SC, SCd: deep SC.

**Figure S5.**
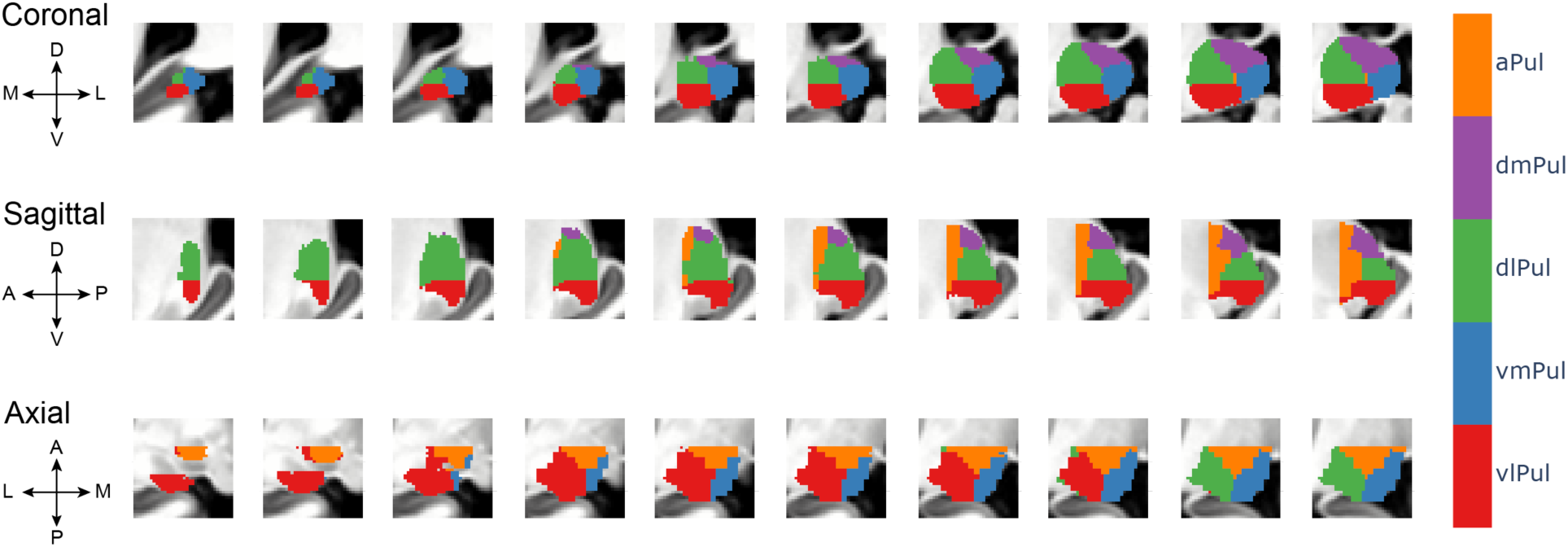
Pulvinar subdivisions. vmPul: ventromedial pulvinar, vlPul: ventrolateral pulvinar, dlPul: dorsolateral pulvinar, dmPul: dorsomedial pulvinar, aPul: anterior pulvinar.

**Figure S6.**
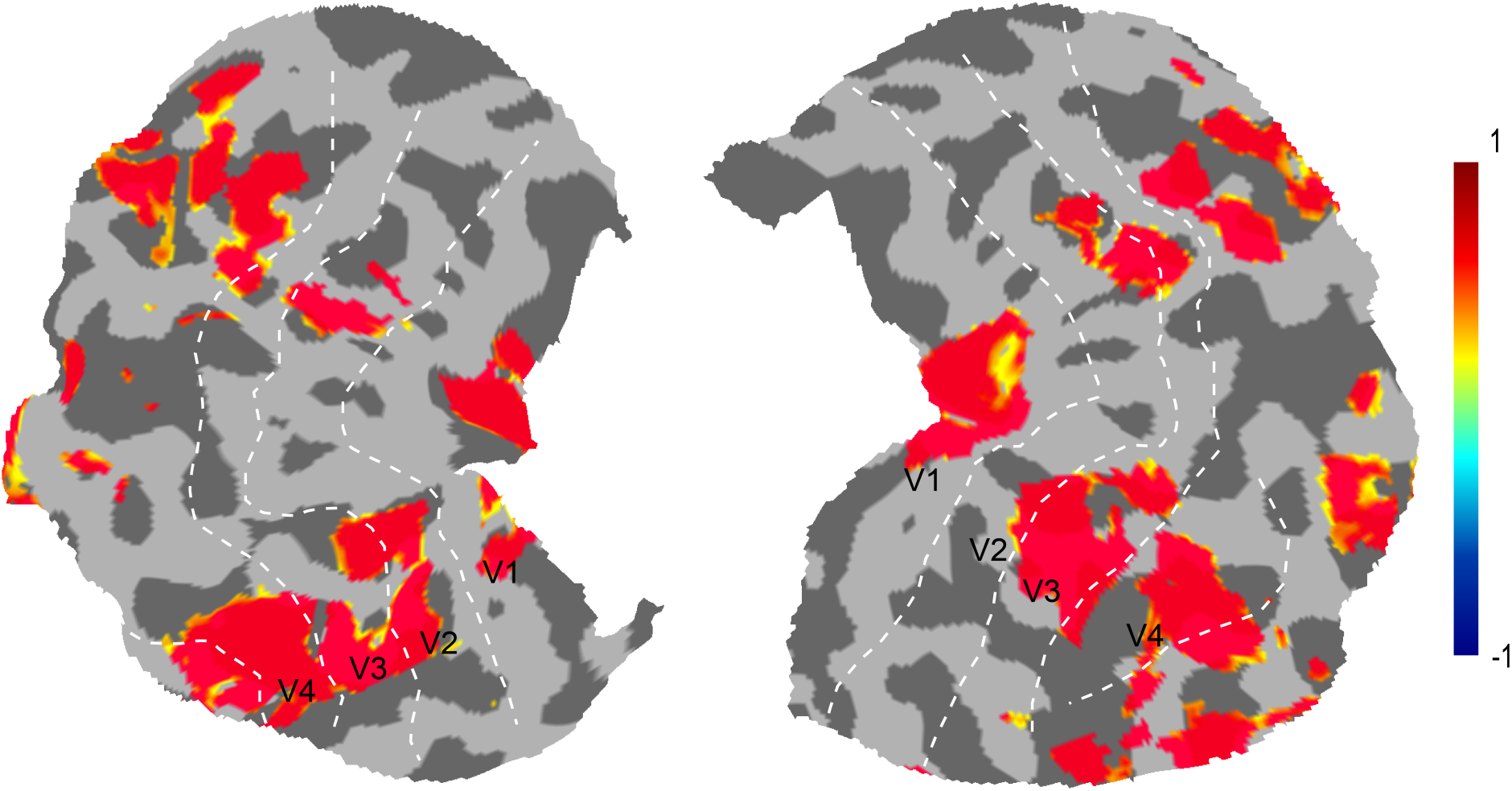
Activations to the chromatic localizer in the early visual cortices in one participant. White dotted lines indicate boundaries of V1-V4 based on the 7T HCP retinotopic atlas. The color bar denotes percent signal change of BOLD response. Activations were thresholded at p < 0.05, uncorrected.

**Figure S7.**
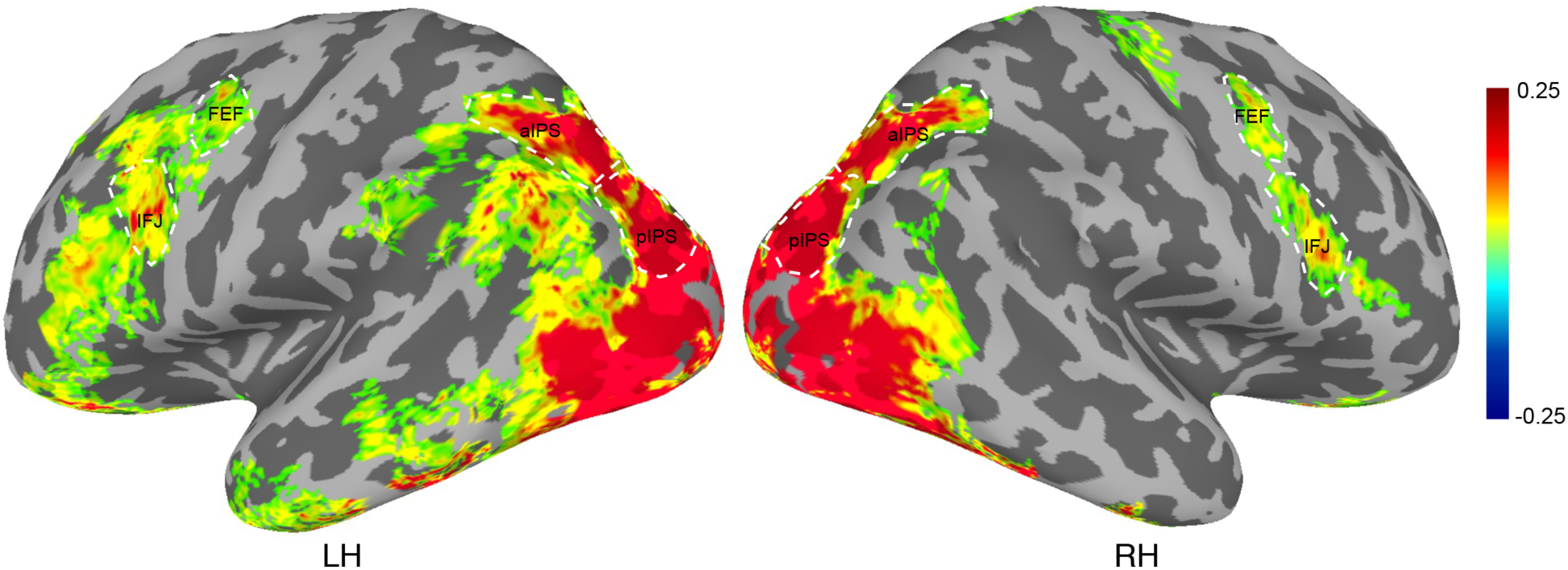
Group-averaged BOLD activation for the chromatic localizer. White dotted lines indicate the boundaries of IFJ, FEF, aIPS and pIPS based on the activation and anatomical landmarks. The color bar denotes percent signal change of BOLD response. Maps were thresholded at t > 0, cluster size less than 300 vertices were not shown.

**Figure S8.**
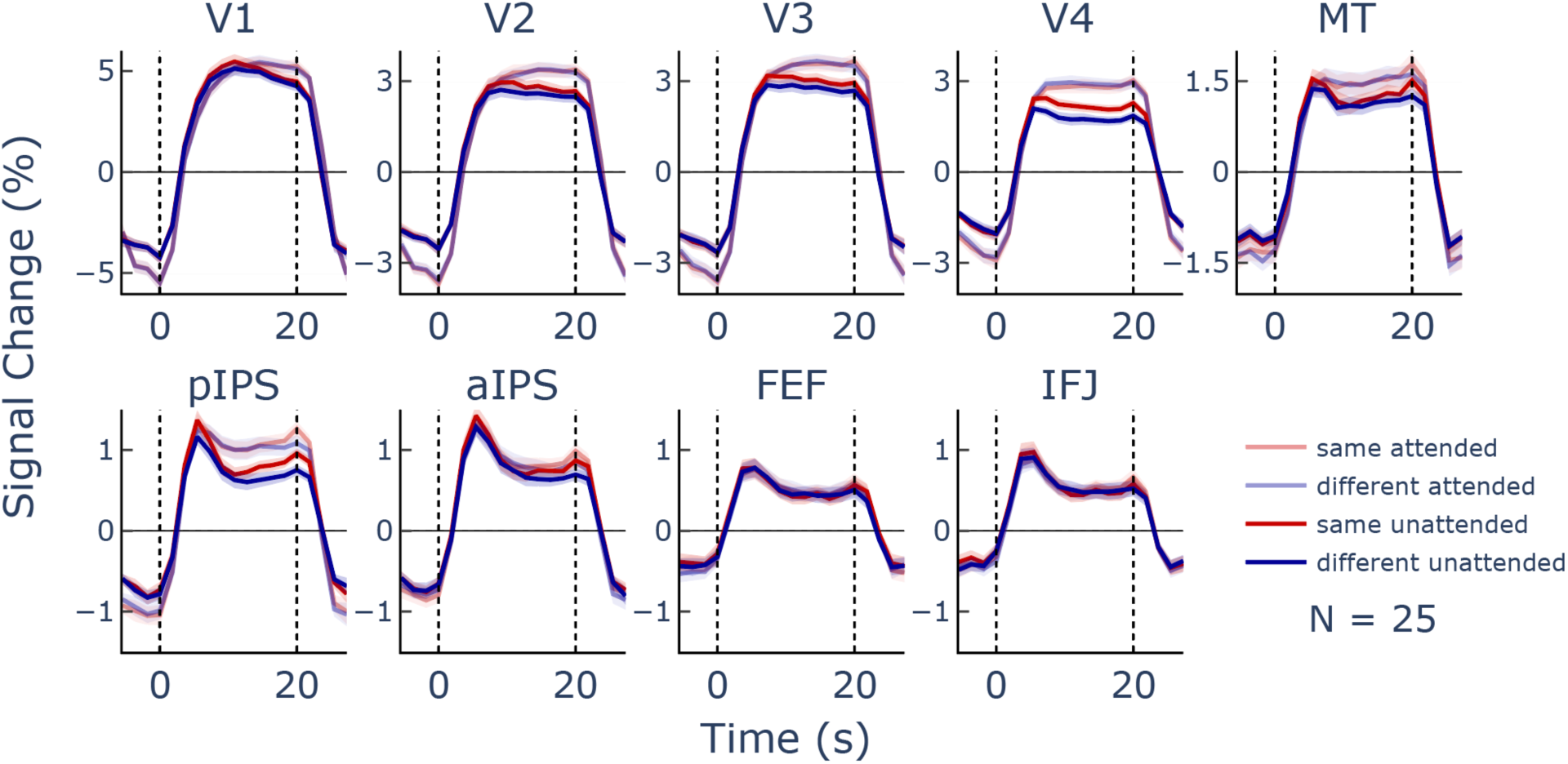
ROI-averaged BOLD timecourses in cortical regions. Shaded areas represent ±SEM.

**Figure S9.**
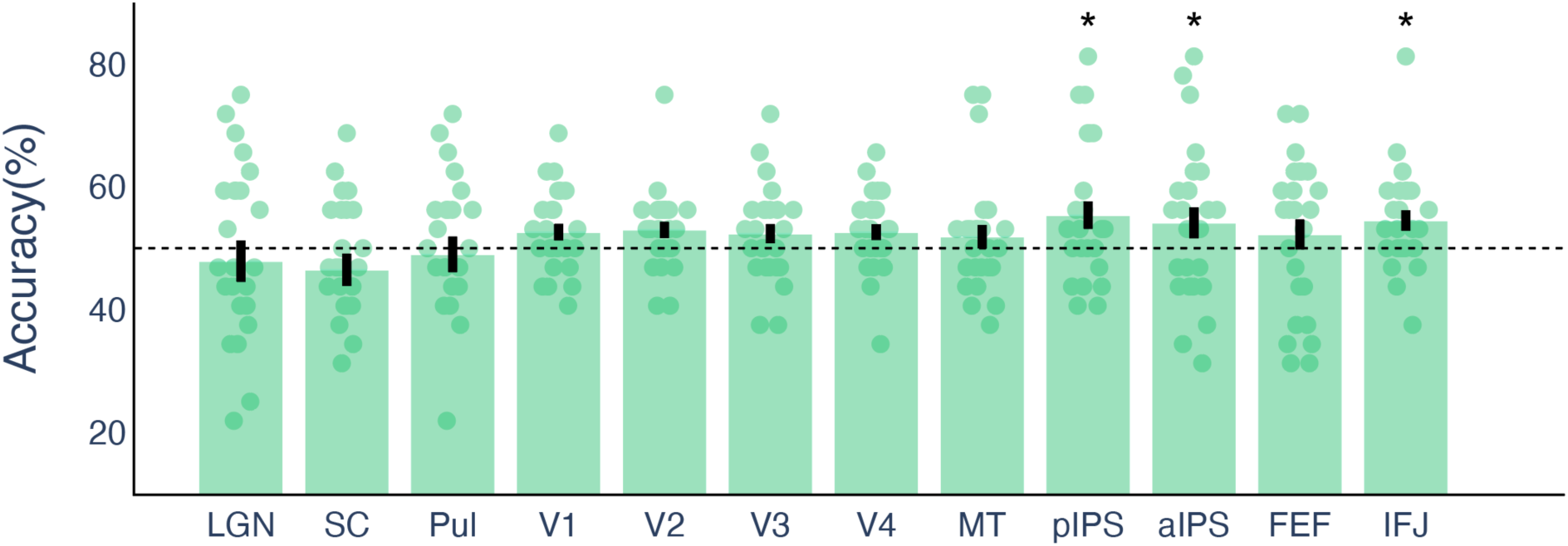
Decoding accuracy of attended color from brain regions contralateral to the attended location in Exp. 2.

**Figure S10.**
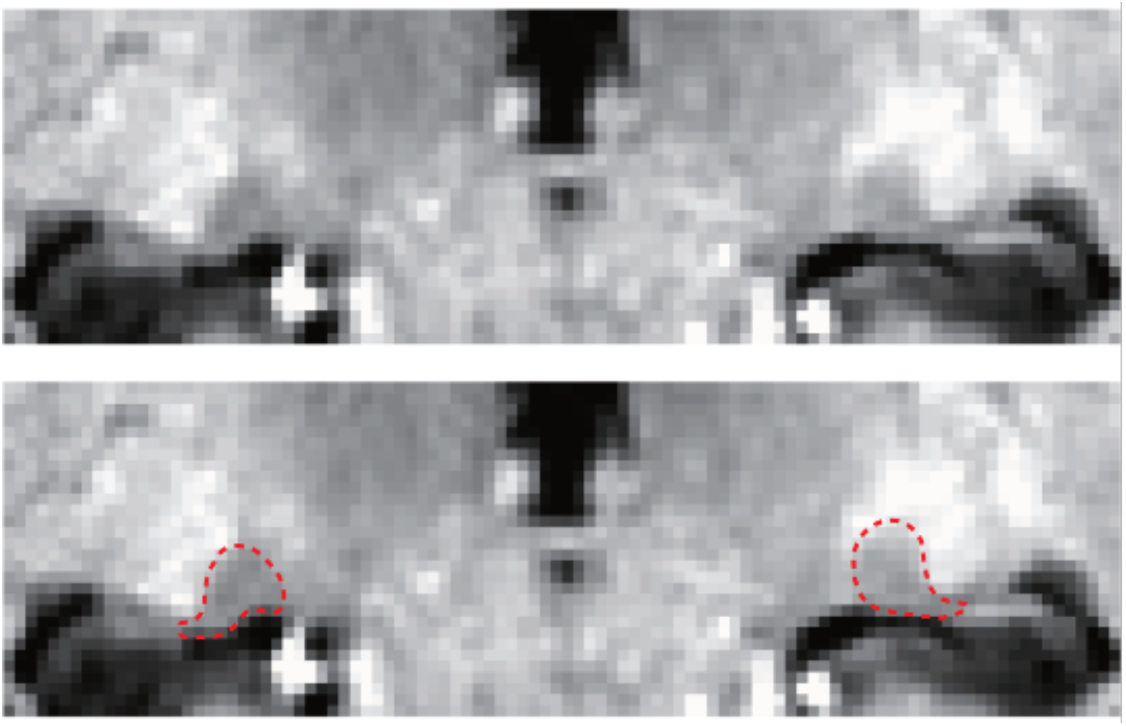
T1-weighted image contrast at 7 T is much improved (due to longer T1 difference and higher image SNR) compared to those at lower magnetic field (Wright et al., 2008). From the T1-weighted MP2RAGE image (0.7-mm isotropic voxels), the anatomical boundary of the LGN is clearly visible thus can be accurately defined by an experienced researcher. The figure below shows the T1-weighted MP2RAGE image around the LGN and the manually defined anatomical boundary (indicated by the red dotted lines). Wright, P. J., Mougin, O. E., Totman, J. J., Peters, A. M., Brookes, M. J., Coxon, R., Morris, P. E., Clemence, M., Francis, S. T., Bowtell, R. W., & Gowland, P. A. (2008). Water proton T1 measurements in brain tissue at 7, 3, and 1.5T using IR-EPI, IR-TSE, and MPRAGE: Results and optimization. Magnetic Resonance Materials in Physics, Biology and Medicine, 21(1–2), 121–130. https://doi.org/10.1007/s10334-008-0104-8

## Notes

### Competing Interest Statement

The authors have declared no competing interest.

